# Neural signatures of opioid-induced risk-taking behavior in the prelimbic prefrontal cortex

**DOI:** 10.1101/2024.02.05.578828

**Authors:** Cana B. Quave, Andres M. Vasquez, Guillermo Aquino-Miranda, Milagros Marín, Esha P. Bora, Chinenye L. Chidomere, Xu O. Zhang, Douglas S. Engelke, Fabricio H. Do-Monte

## Abstract

Opioid use disorder occurs alongside impaired risk-related decision-making, but the underlying neural correlates are unclear. We developed an approach-avoidance conflict task using a modified conditioned place preference procedure to study neural signals of risky opioid seeking in the prefrontal cortex, a region implicated in executive decision-making. Following morphine conditioned place preference, rats underwent a conflict test in which fear-inducing cat odor was introduced in the previously drug-paired side of the apparatus. While the saline-exposed control group avoided cat odor, the morphine group included two subsets of rats that either maintained a preference for the paired side despite the presence of cat odor (Risk-Takers) or exhibited increased avoidance (Risk-Avoiders), as revealed by K-means clustering. Single-unit recordings from the prelimbic cortex (PL) demonstrated decreased neuronal activity upon acute morphine exposure in both Risk-Takers and Risk-Avoiders, but this firing rate suppression was absent after repeated morphine administration. Risk-Avoiders also displayed distinct post-morphine excitation in PL which persisted across conditioning. During the preference test, subpopulations of PL neurons in all groups were either excited or inhibited when rats entered the paired side. Interestingly, the inhibition in PL activity was lost during the subsequent conflict test in both saline and Risk-Avoider groups, but persisted in Risk-Takers. Additionally, Risk-Takers showed an increase in the proportion of PL neurons displaying location-specific firing in the drug-paired side from the preference to the conflict test. Together, our results suggest that persistent PL inhibitory signaling in the drug-associated context during motivational conflict may underlie increased risk-taking behavior following opioid exposure.

**SIGNIFICANCE STATEMENT:** Risky opioid use is well established in opioid use disorder, but the underlying neural correlates are poorly understood. In this study, we present findings from a novel behavioral task in which rats face a motivational conflict between contextual opioid reward memory and a naturalistic predator threat. Performing neuronal recordings in the prelimbic prefrontal cortex (PL), a brain region critical for executive decision-making, we demonstrate enhanced representation of drug-associated context and persistent inhibitory signaling by PL neurons that occur alongside opioid-induced risk-taking behavior. Our findings refine a preclinical model for studying addiction, establish PL as a prime region for investigating drug-environment interactions, and positions the prefrontal cortex as a candidate region for translational studies targeting risky opioid use.

## INTRODUCTION

Opioid use disorder is associated with deficits in risk-related decision-making (Petry et al., 1998; Barry and Petry, 2008). While much has been done to elucidate the biological mechanisms of reward seeking during neutral conditions (Wise, 2002; Berridge, 2007), relatively little is known about drug seeking in risky circumstances. Prior studies have sought to model risky opioid self-administration in laboratory animals, and results have shown that rodents will continue to seek opioids even when they must risk experiencing painful stimuli (e.g., electrical footshocks) to do so (Panlilio et al., 2003, 2005; Porter-Stransky et al., 2017; Blackwood et al., 2020; Borges et al., 2022, 2023; Honeycutt et al., 2022; Mathieson et al., 2022). Other studies have further attempted to model opioid use in a risky context by introducing an electrified barrier that animals had to cross to obtain the drug (Peck et al., 2013, 2015; Fredriksson et al., 2020, 2021a, 2023; Martin et al., 2022; Negishi et al., 2024). However, experimental use of painful stimuli as impediments to drug seeking does not fully resemble the harms associated with obtaining or using drugs in humans (De Peuter et al., 2011). The use of nociceptive stimuli, such as footshock, to investigate risky opioid seeking is further complicated by the fact that repeated opioid use can alter nociceptive sensitivity (Liebmann et al., 1997; Lee et al., 2011), thereby resulting in confounding findings.

One animal model that has been extensively used to study opioid reward is the conditioned place preference (CPP) paradigm (Mucha et al., 1982; Cunningham et al., 2006). Although CPP limits interpretations regarding drug *taking*, some motivational aspects of drug *seeking* are present in that the animal must “choose” to enter and remain in a drug-paired context (Green and Bardo, 2020). With its ability to capture motivated behaviors related to drug-reward memory in the absence of the drug, CPP is an ideal model for studying drug preference as a function of environmental context. Thus, to define the phenotype of opioid-induced risk-taking behavior in rats, we developed an approach-avoidance conflict model that pits contextual drug memory against predator odor-induced fear by exposing rats to CPP conditioning with opioids followed by the introduction of a non-nociceptive aversive stimulus (*i.e.*, cat saliva; Papes et al., 2010; Engelke et al., 2021) in the drug-associated context.

The medial prefrontal cortex (mPFC) is critical for top-down cognitive control of emotionally motivated behaviors in humans, but its role in risky reward seeking has not been fully characterized (Goldstein and Volkow, 2011). In rodents, neurons in the prelimbic subregion (PL) of the mPFC are activated by cues that predict either rewarding or threatening stimuli (Burgos-Robles et al., 2009; Sierra-Mercado et al., 2011; Sangha et al., 2014; Do-Monte et al., 2015; Otis et al., 2017; Fernandez-Leon et al., 2021). Additionally, aberrant activity in PL has been implicated in the dysregulation of goal-directed behavior and the persistence of drug seeking despite aversive consequences (Kasanetz et al., 2013; Smith and Laiks, 2018; Hu et al., 2019), making this region a potential candidate to regulate risky decision-making. Of particular relevance to opioid seeking, activation of opioid receptors in PL neurons is necessary for the formation of contextual opioid reward memory (Jiang et al., 2021), and the expression of this opioid-associated memory is blocked by chemogenetic silencing of PL (Hou et al., 2018).

We therefore hypothesized that the signaling of drug-related contextual information in PL is suppressed in the presence of threat, and that failure of this suppression occurs during risky opioid seeking. To further investigate the prefrontal cortex mechanisms that underlie opioid-induced risky decision making, we performed *in vivo* electrophysiological recordings from PL neurons in freely moving rats to assess changes in PL activity during the development of opioid CPP, as well as to identify patterns of contextual representation in PL neurons during the opioid-approach versus predator threat-avoidance conflict test. Our results reveal prefrontal neural correlates of opioid-induced risk-taking in a drug-associated context.

## MATERIALS AND METHODS

### Animals

Adult male and female Long-Evans hooded rats (Charles River Laboratories) were used. All rats were 3-5 months of age and weighed ∼350 to 500 g at the time of testing. Rats were maintained on a 12-h light / 12-h dark cycle (7:00 to 19:00 light period) with *ad libitum* access to water. Rats were also maintained on a restricted diet of standard laboratory rat chow (18 g per day) and weighed weekly to ensure all rats maintained their weights throughout the course of experimentation. All experiments were approved by The University of Texas Health Science Center at Houston Center for Laboratory Animal Medicine and Care. The National Institutes of Health Guide for the Care and Use of Laboratory Animals was followed in order to prevent unnecessary animal suffering or discomfort.

### Drugs

All drugs used in behavioral experiments were injected subcutaneously. Only pharmaceutical-grade morphine sulphate (10 mg/mL, Hikma) or fentanyl citrate (2,500 mcg/50mL, West-Ward) prepared for human intravenous use were administered to rats during the course of the study.

More information about all experimental methods including cat odor collection and preparation, behavioral tasks, stereotaxic surgeries, in vivo single-unit electrophysiology, histology, statistics and reproducibility are presented in detail in the **Supplementary Methods** section.

### Opioid-approach versus predator threat-avoidance conflict model

Rats underwent conditioning in a two-chamber apparatus. On Day 1, rats freely explored both sides of the apparatus and baseline side preferences were recorded. The following day, rats were injected with either saline, morphine, or fentanyl and confined to the side of the apparatus preferred least at baseline. Conditioning occurred over 10 alternating days (5 pairings in each side). On day 13, rats underwent a 10-min preference test immediately followed by a

10-min conflict test in which an aversive stimulus (cat saliva) was introduced in the side of the chamber previously paired with drug injections and side preference/aversion were again recorded.

### Experimental design and statistical analysis

Shapiro-Wilk normality tests were first used to assess normality. For group behavioral data, one-way or two-way ANOVA were used as appropriate, followed by Šidák’s multiple comparisons tests to investigate significant interactions. For comparison of behavioral data between two groups or area under the curves (AUC) for single-unit data, we employed either unpaired Welch’s t-tests or paired Student’s t-tests for normally-distributed data, and Mann-Whitney *U*-tests or Wilcoxon matched-pairs signed rank tests for non-normally distributed unpaired or paired data, respectively. For the comparison of categorical variables, either Fisher’s Exact test or the Chi-square test was used. Sample sizes were determined using power analysis, with a significance level of 0.05 and a power of 0.8.

## RESULTS

### Morphine conditioning leads to contextual place preference and individual differences in risk-taking behavior in rats

To study opioid-induced risk-taking behavior, we modified a traditional CPP protocol to include a component of approach-avoidance conflict. First, adult male rats were assigned to one side of a two-chamber apparatus for conditioning, with the assigned side being the one they least preferred at baseline. Rats were then conditioned with systemic administration of either morphine (10 mg/kg, s.c.) or saline (1 mL/kg) in alternating sides every other day for 10 days. All rats received five drug pairings in one side of the apparatus (drug-paired side) and five saline pairings in the other (neutral side; **Fig. 1A**). A control group (saline-treated rats) were subjected to an identical conditioning protocol, but received saline injections alternately in either side of the apparatus on all 10 conditioning days. During the preference test performed two days after conditioning, morphine-treated rats showed CPP as evidenced by increased time spent in the apparatus’ drug-paired side (**Fig. 1B**, Welch’s t-test, *p* = 0.0004; **Supplementary Fig. 1**, two-way repeated measures ANOVA (F _(2, 114)_ = 4.990, Day X Drug interaction, *p* = 0.0084, Šidák’s multiple comparisons test, baseline vs. preference test, *p* = 0.0004). Immediately following the preference test, the rats were briefly removed from the apparatus and an aversive stimulus (cat saliva) was placed in the drug-paired side (see details in the Methods section). Cat saliva has been shown to elicit innate defensive behaviors in rodents, including avoidance of the cat odor source (Papes et al., 2010; Engelke et al., 2021).

**Figure 1.**
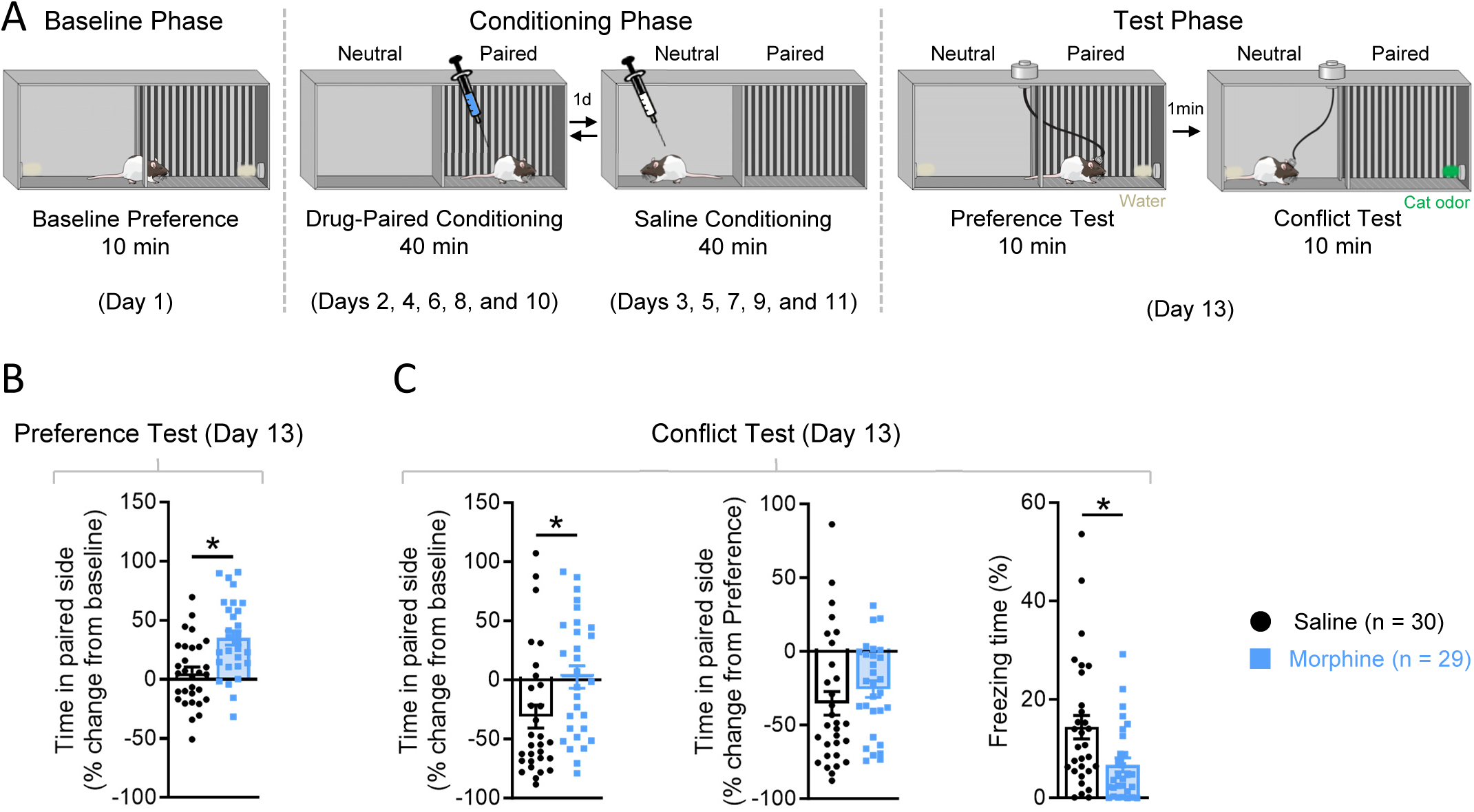
Repeated morphine administration leads to contextual reward memory formation and risk-taking behavior during approach-avoidance conflict. **A)** Schematic timeline of morphine conditioned place preference and approach-avoidance conflict tests. Rats were assigned to one side of a two-chamber apparatus for conditioning, the assigned side being that which the rat preferred least at baseline. **B)** Percentage of change from baseline in time spent in the drug-paired side of the apparatus. Morphine-treated rats exhibited conditioned place preference as measured by the increased amount of time in the drug-paired side compared to saline controls (Welch’s t-test, \**p* < 0.001). **C *left*)** Percentage of change from baseline in time spent in the drug/cat-paired side of the apparatus during the conflict test. Rats conditioned with saline, but not morphine, showed aversion to the drug/cat-paired side (Shapiro-Wilk normality test, *p* < 0.001; Mann-Whitney *U*-test, \**p* = 0.0062). **C *center*)** Percentage of change from the preference test in time spent in the drug/cat-paired side of the apparatus during the conflict test. No groups differences were observed (Shapiro-Wilk normality test, *p* = 0.007; Mann-Whitney *U*-test, *p* = 0.149). **C *right*)** Percentage of time spent freezing during the conflict test. Morphine-treated rats displayed reduced freezing levels compared to saline controls (Shapiro-Wilk normality test, *p* < 0.05; Mann-Whitney *U*-test, \**p* = 0.0053). Data are shown as mean ± SEM.

After the addition of cat odor, rats were then returned to the apparatus for a motivational conflict test in which the rats had to approach cat odor to visit the drug-associated chamber. We observed that saline-treated rats avoided the side of the apparatus containing cat odor (**Fig. 1C *left***, Mann-Whitney *U*-test, *p* = 0.0062; **Supplementary Fig. 1**, two-way repeated measures ANOVA (F _(2, 114)_ = 4.990, Day X Drug interaction, *p* = 0.0084, Šidák’s multiple comparisons test, preference test vs. conflict test, *p* < 0.0001). In contrast, some opioid-treated rats continued to enter the drug-paired side despite the presence of cat odor, suggesting that morphine conditioning increases risk-taking behavior (**Fig. 1C *left***, Mann-Whitney *U*-test, *p* = 0.0062, **1C *center***; **Supplementary Fig. 1**, two-way repeated measures ANOVA (F _(2, 114)_ = 4.990, Day X Drug interaction, *p* = 0.0084, Šidák’s multiple comparisons test, saline vs. morphine, *p* = 0.0014). Moreover, morphine-treated rats displayed reduced defensive responses as measured by lower levels of freezing behavior as compared to saline controls (**Fig. 1C *right***, Mann-Whitney *U*-test, \**p* = 0.0053).

### Repeated morphine administration in the home cage does not affect defensive behavior

Because morphine-treated rats exhibited decreased freezing levels during the conflict test, we questioned if opioid exposure leads to a reduction in general defensive responses to cat odor due to impaired olfactory function and/or reduced antipredator defense. To test this, we performed a new experiment using the same dosing schedule as before, but instead of conditioning the rats in the apparatus, we returned them to their home cages following morphine injections (**Supplementary Fig. 2A**). During the cat odor test performed two days after the last injection, morphine- and saline-treated rats spent similar amounts of time in the side of the apparatus containing cat odor (**Supplementary Fig. 2B *left***, Welch’s t-test, *p* = 0.27). We observed that cat odor evoked increased freezing behavior in both saline- and morphine-treated rats compared to baseline (two-way repeated measures ANOVA, Cat Odor main effect, F _(1, 18)_ = 33.67, *p <* 0.0001, Šidák’s multiple comparisons tests, saline: *p* = 0.0048, morphine: *p* = 0.0004), and the two groups exhibited similar defensive responses to cat odor as measured by either freezing (**Supplementary Fig. 2B *center***, Welch’s t-test, *p* = 0.528) or total distance traveled in the apparatus (**Supplementary Fig. 2B *right***, Welch’s t-test, *p* = 0.318) during the test. These results suggest that repeated exposure to morphine, in the dose and schedule used in our main experiment, does not affect general defensive responses to cat odor outside of a drug-associated context. Importantly, these results support the conclusion that risk-taking behavior as observed in our main experiment likely depends on the contextual association of the apparatus with the rewarding effects of morphine, rather than simply on the pharmacological effects of the drug.

### High doses of opioids are necessary to induce risk-taking behavior

The rewarding effects of morphine are dose-dependent (Bardo et al., 1995). To test whether conditioning with a lower dose of morphine can produce risk-taking behavior in our model, we repeated the same experiments using injections of 5 mg/kg, half of our previous dose. This dose of morphine was insufficient to produce either CPP during the preference test (**Supplementary Fig. 3A, M**ann-Whitney *U*-test, *p* = 0.336; **Supplementary Fig. 3E**, two-way repeated measures ANOVA, Test main effect, F _(2, 84)_ = 9.13, *p* < 0.0001, no Drug x Test interaction, F _(2, 84)_ = 0.66, *p* = 0.52) or risk-taking behavior during the conflict test (**Supplementary Fig. 3B, M**ann-Whitney *U*-test, *p* = 0.54; **Supplementary Fig. 3E**, two-way repeated measures ANOVA, Test main effect, F _(2, 84)_ = 9.13, *p* < 0.0001, no Drug x Test interaction, F _(2, 84)_ = 0.66, *p* = 0.52). These results reveal that higher doses of morphine are required to elicit both morphine preference and risk-taking behavior under our experimental conditions.

Recent reports have shown that fentanyl, a synthetic opioid drug ∼100 times more potent than morphine, contributes largely to overdose-related deaths in humans (Scholl, 2019). We sought to determine if fentanyl can induce risk-taking behavior in our model in a manner similar to that observed with morphine. As with morphine, we performed experiments using two separate doses of fentanyl (0.02 or 0.04 mg/kg). Both fentanyl doses led to CPP during the preference test (**Supplementary Fig. 3C, 0**.02 mg/kg: Welch’s t-test, *p* = 0.0003, 0.04 mg/kg: Welch’s t-test, *p* = 0.0005; **Supplementary Fig. 3E**). However, as we have observed with morphine administration, only rats treated with the higher dose of fentanyl showed risk-taking behavior during conflict (**Supplementary Fig. 3D, 0**.02 mg/kg: Mann-Whitney test, *p* = 0.14, 0.04 mg/kg: Mann-Whitney *U*-test, *p* = 0.0396; **Supplementary Fig. 3E**), reinforcing our observation that higher doses of opioids are necessary to promote risky behavior. These results also indicate that although a drug’s ability to induce contextual reward memory is associated with its potential to affect risk-related decision making, these effects appear to be separable.

### Male rats show greater sensitivity than female rats to opioid-induced risk-taking behavior at equivalent doses

Although sex differences in opioid addiction-related behaviors have been previously reported (Chartoff and McHugh, 2016; Knouse and Briand, 2021), the broad interpretation of individual findings across specific behavioral domains has recently been called into question (Nicolas et al., 2022). Additionally, research investigating sex differences in opioid seeking despite negative consequences is scarce, and the few existing results show similar behavior or sex-dependent differences according to the test (Fredriksson et al., 2020; Monroe and Radke, 2021). To investigate sex differences in risk-taking behavior in our model, we performed the same experiments (described above) in adult female rats using the highest doses of either morphine or fentanyl that produced CPP and risk-taking behavior in males.

We found that during the preference test, females that were injected with fentanyl, but not those injected with morphine, showed CPP (**Supplementary Fig. 4A *left***, Welch’s t-test, *p* = 0.64, **4A *right***, Mann-Whitney *U*-test, *p* = 0.0011; **Supplementary Fig. 4C**). Furthermore, fentanyl-conditioned females displayed increased risk-taking behavior during the conflict test, while those conditioned with morphine did not (**Supplementary Fig. 4B *left***, Mann-Whitney *U*-test, *p* = 0.37, **4B *right***, Mann-Whitney *U*-test, *p* = 0.0031; **Supplementary Fig. 4C**). Therefore, we conducted further experiments solely with males using the highest dose of morphine (10 mg/kg), which produced consistent behavioral effects in this group of rats.

### Risk-Taker and Risk-Avoider behavioral phenotypes emerge in an opioid-approach versus predator threat-avoidance conflict test

Studies of risky drug seeking using other conflict models have identified two distinct behavioral phenotypes in drug-exposed rodents: i) those that ceased drug taking when confronted with risk of shock punishment, and ii) those that were aversion-resistant and continued to pursue drugs despite risk (Deroche-Gamonet et al., 2004; Chen et al., 2013; Porter-Stransky et al., 2017; Marchant et al., 2018; Venniro et al., 2018; Blackwood et al., 2020). To determine if differential risk-taking phenotypes also exist within our model, we first identified behaviors we considered relevant to risk taking, including CPP (**Fig. 1B**), cat odor aversion (reduction in time spent in the paired side from the preference test to the conflict test; **Fig. 1C *center***), and freezing (**Fig. 1C *right***). It is important to note that the first two measures, CPP during the preference test and cat odor aversion during the conflict test, reflect conceptually separate phenomena: CPP indicates side preference, whereas cat odor aversion reflects an independent response to a predator stimulus, as demonstrated by our previously described control experiment (**Supplementary Fig. 2**). We then performed a K-means clustering of data from these three measures, which revealed two subgroups of morphine-treated rats. The first subgroup showed enhanced CPP and increased risk-taking behavior during conflict (Risk-Takers), whereas the second subgroup exhibited suppressed CPP and pronounced cat odor avoidance during conflict (Risk-Avoiders; **Fig. 2A**). Importantly, when we plotted data from saline-treated rats alongside data from Risk-Avoiders and Risk-Takers on these three dimensions, we observed a 40% (12 of 30 rats) overlap of the saline-treated group cluster with the Risk-Avoider group cluster, in contrast to only 3% (1 of 30 rats) overlap with the Risk-Taker group cluster (**Supplementary Fig. 5A, F**isher’s Exact test, ratios of overlapping to non-overlapping rats, *p* = 0.0011; see Methods section). Compared to Risk-Avoiders, Risk-Takers demonstrated greater measures of CPP during both the preference and conflict tests (**Fig. 2B**, Welch’s t-test, *p* = 0.004; **Fig. 2C *left***, Welch’s t-test, *p* < 0.0001; **Supplementary Fig. 5B–C**; one-way ANOVA, F _(2, 56)_ = 16.69, *p* < 0.0001, Šidák’s multiple comparisons test, *p* = 0.0008; **Supplementary Fig 5D *left***, one-way ANOVA, F _(2, 56)_ = 20.67, *p* < 0.0001, Šidák’s multiple comparisons test, *p* < 0.0001), as well as a lack of aversion to cat odor during the conflict test (**Fig. 2C *center***, Welch’s t-test, \**p* < 0.0001; **Fig. 2D–E**; **Supplementary Fig. 5B**, two-way repeated measures ANOVA, Group x Test interaction, F _(2, 54)_ 37.93, *p <* 0.0001, Šidák’s multiple comparisons test, *p* < 0.0001; **Supplementary Fig. 5D *center***, one-way ANOVA, F _(2, 56)_ = 6.302, *p* = 0.0034, Šidák’s multiple comparisons test, *p* = 0.0039). Compared to saline-treated rats, Risk-Takers similarly showed CPP during the preference and conflict tests (**Supplementary Fig. 5C**, one-way ANOVA, F _(2, 56)_ = 16.69, *p* < 0.0001, Šidák’s multiple comparisons test, *p* < 0.0001; **Supplementary Fig. 5D *left***, one-way ANOVA, F _(2, 56)_ = 20.67, *p* < 0.0001, Šidák’s multiple comparisons test, *p* < 0.0001), and reduced cat odor aversion (**Supplementary Fig. 5D *center***, one-way ANOVA, F _(2, 56)_ = 6.302, *p* = 0.0034, Šidák’s multiple comparisons test, *p* = 0.031). In contrast, Risk-Avoiders did not differ from saline-treated rats on measures of CPP in the preference or conflict tests (**Supplementary Fig. 5C**, one-way ANOVA, F _(2, 56)_ = 16.69, *p* < 0.0001, Šidák’s multiple comparisons test, *p* = 0.62; **Supplementary Fig. 5D *left***, one-way ANOVA, F _(2, 56)_ = 20.67, *p* < 0.0001, Šidák’s multiple comparisons test, *p* = 0.75), nor cat odor aversion (**Supplementary Fig. 5D *center***, one-way ANOVA, F _(2, 56)_ = 6.302, *p* = 0.0034, Šidák’s multiple comparisons test, *p* = 0.59). Notably, while only Risk-Takers exhibited reduced freezing during the conflict test compared to saline-treated rats (**Supplementary Fig. 5D *right***, one-way ANOVA, F _(2, 56)_ = 4.9, *p* = 0.011, Šidák’s multiple comparisons test, saline vs. Risk-Avoiders: *p* = 0.44, saline vs. Risk-Takers: *p* = 0.0089), Risk-Avoiders and Risk-Takers did not differ in levels of freezing (**Fig 2C *right***, Welch’s t-test, *p* = 0.052; **Supplementary Fig. 5D *right***, one-way ANOVA, F _(2, 56)_ = 4.9, *p* = 0.011, Šidák’s multiple comparisons test, *p* = 0.39), further suggesting that opioid-induced risk-taking behavior is not due to a general suppression of defensive behavior.

**Figure 2.**
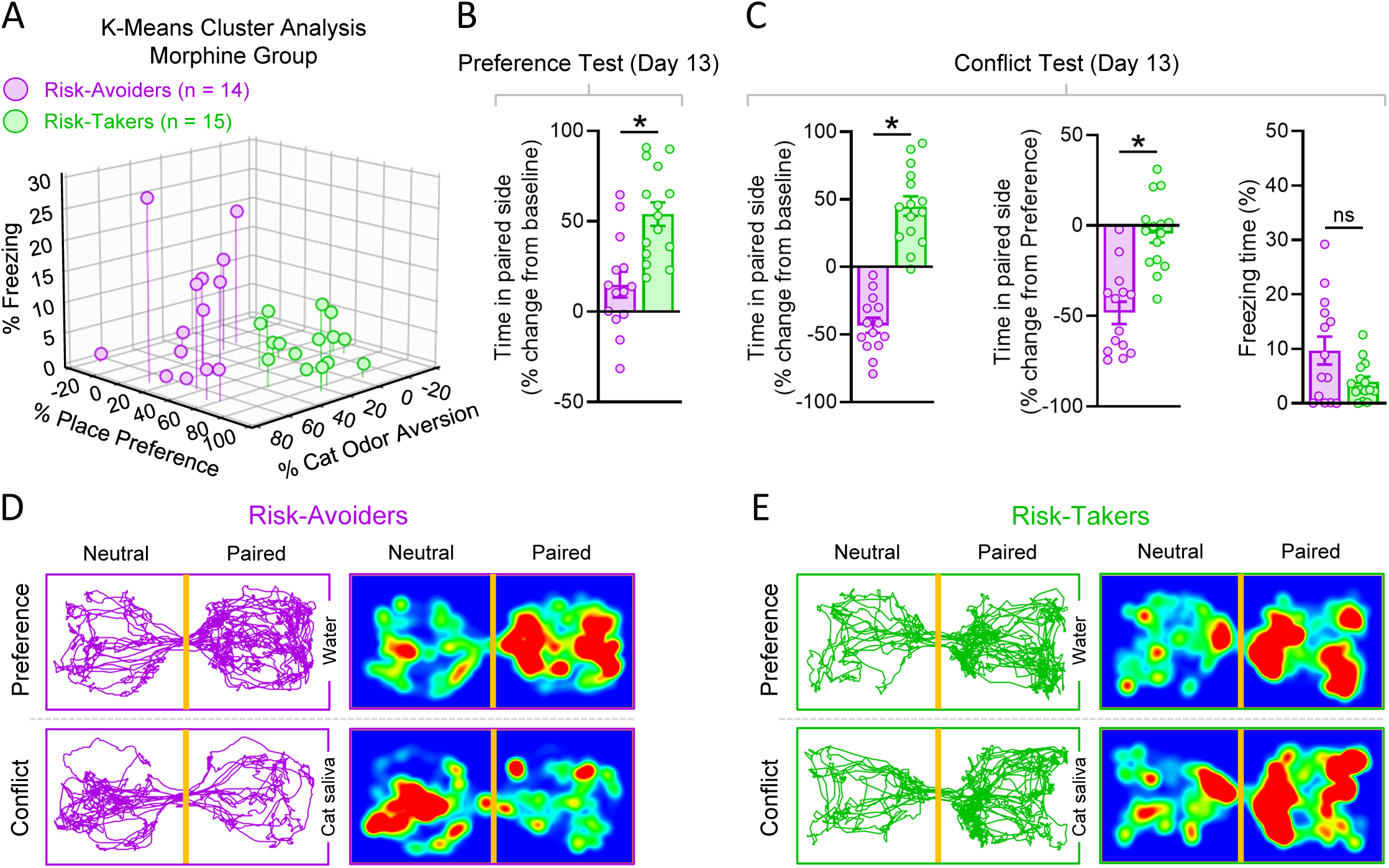
Morphine-treated rats show individual differences in risk-taking behavior during conflict. **A)** K means cluster analysis (10 repetitions) of Morphine group animals in measures of freezing (% time spent freezing during the conflict test), place preference (% change from baseline in time spent in the drug-paired side during the preference test), and cat odor aversion (% change from preference test in time spent in the drug/cat-paired side during the conflict test). Two clusters were identified: one with lower place preference and greater cat odor aversion (magenta cluster, Risk-Avoiders, n = 15), and another with greater place preference and lower cat odor aversion (green cluster, Risk-Takers, n = 14). **B)** Percentage of change from baseline in time spent in the drug-paired side of the apparatus. Risk-Takers demonstrated greater place preference than Risk-Avoiders (Welch’s t-test, \**p* < 0.001). **C *left*)** Percentage of change from baseline in time spent in the drug/cat-paired side of the apparatus during the conflict test. Risk-Takers showed less cat odor aversion than Risk-Avoiders (Welch’s t-test, \**p* < 0.0001). **C *center*)** Percentage of change from the preference test in time spent in the drug/cat-paired side of the apparatus during the conflict test. Risk-Takers showed less cat odor aversion than Risk-Avoiders (Welch’s t-test, \**p* < 0.0001). **C *right*)** Percentage of time spent freezing during the conflict test. Both groups displayed similar levels of freezing during the conflict test (Welch’s t-test, \**p* = 0.052). Representative tracks and heatmaps of time spent in either side of the apparatus during Preference or Conflict Tests for Risk-Avoiders **(D)** and Risk-Takers **(E)**. Data are shown as mean ± SEM.

### Risk-taking behavior cannot be explained by individual differences in CPP strength or extinction

While persistent CPP despite the presence of a threat can be interpreted as risk-taking behavior, alternative interpretations are also possible. One such interpretation is that variability in the strength of morphine conditioning can lead to differences in CPP expression that correlate with risk-taking behavior during the conflict test. To test this, we performed K-means clustering of CPP data from morphine-treated rats during the preference test to identify low-preference and high-preference groups (**Supplementary Fig. 6A *left***, one-way ANOVA, F _(2, 56)_ = 29.96, *p* < 0.0001, Šidák’s multiple comparisons tests, saline vs. low-preference: *p* = 0.80, saline vs. high-preference: *p* < 0.0001, low-preference vs. high-preference: *p* < 0.0001). In contrast to Risk-Avoiders and Risk-Takers, which differed in expression of CPP during the conflict test (**Fig. 2C *left***, Welch’s t-test, *p* < 0.0001; **Supplementary Fig. 5D *left***, one-way ANOVA, F _(2, 56)_ = 20.67, *p* < 0.0001, Šidák’s multiple comparisons test, *p* < 0.0001), low-preference and high-preference rats displayed similar percentages of time spent in the drug-paired side during the conflict test (**Supplementary Fig. 6A *center***, one-way ANOVA, F _(2, 56)_ = 5.968, *p* = 0.0045, Šidák’s multiple comparisons test, *p* = 0.0697). Importantly, high-preference rats did not show reduced cat odor aversion during the conflict test (**Supplementary Fig. 6A *right***, one-way ANOVA, F _(2, 56)_ = 0.5207, *p* = 0.597, Šidák’s multiple comparisons test, saline vs. high-preference: *p* = 0.72), as was apparent in Risk-Takers (**Fig. 2C *center***, Welch’s t-test, \**p* < 0.0001; **Supplementary Fig. 5D *center***, one-way ANOVA, F _(2, 56)_ = 6.302, *p* = 0.0034, Šidák’s multiple comparisons test, *p* = 0.031). These data indicate that CPP expressed during the preference test and risk taking during the conflict test are not mutually dependent; this point is further evidenced by the observation that 33% of the Risk-Taker group was comprised of low-preference rats (**Supplementary Fig. 6B**), whereas 31% of the low-preference group was comprised of Risk-Takers (**Supplementary Fig. 6C**). Together, these results suggest that while enhanced CPP might be important for the expression of risk-taking behavior, it is certainly not required.

A second possible interpretation of our results is that Risk-Avoiders and Risk-Takers differ during the conflict test because of distinct rates of CPP extinction during the preference test. However, after segregating CPP data from the 10-min preference test into sequential 2-min bins, we found that time spent in the drug-paired side did not diminish across the duration of the preference test in any subgroup of rats (**Supplementary Fig. 6D *left***, two-way ANOVA, F _(4, 228)_ = 1.414, *p* = 0.23; **Supplementary Fig. 6D *right***, two-way ANOVA, F _(8, 224)_ = 0.8224, *p* = 0.584), ruling out the possibility that behavioral differences between Risk-Avoiders and Risk-Takers during the conflict test are due to differences in CPP extinction during the preference test. Thus, differences in risk-taking behavior in our test seems to be mostly related to differences in motivational conflict rather than merely the strength or maintenance of opioid conditioning.

Some of the previous studies that identified individual risk-taking phenotypes in drug-seeking animals found that risk taking was associated with other addiction-like behaviors, including increased drug intake, higher progressive ratio breakpoints, and drug-induced reinstatement of seeking responses (Deroche-Gamonet et al., 2004; Chen et al., 2013; Porter-Stransky et al., 2017; Venniro et al., 2018; Blackwood et al., 2020). To determine if risk taking in our model was similarly associated with separate behaviors related to drug-reward, we performed correlation analyses between behavioral measures obtained during the conflict test (paired-side preference or aversion) and those obtained earlier in the experiment (during the preference test). We found no correlation between paired-side preference during the preference and conflict tests in Risk-Avoiders, Risk-Takers, or saline-treated controls. However, we did find that paired-side preference during the conflict test was correlated with increased CPP during the preference test when both groups of morphine-treated rats were combined (**Supplementary Fig. 7A–D**, saline: *r*(28) = 0.33, *p* = 0.08, Risk-Avoiders: *r*(12) = -0.11, *p* = 0.71, Risk-Takers: *r*(13) = 0.32, *p* = 0.25, combined morphine: *r*(27) = 0.59, *p* < 0.001). This significant correlation between behaviors in the preference and conflict tests observed exclusively in the morphine-treated group suggests that the ability of morphine to induce risk-taking behavior in our model is strongly linked to its capacity to elicit contextual reward memory at the individual level. This idea aligns with our findings from correlation analyses of female data showing lack of association between CPP and risk-taking behavior at a non-rewarding dose of morphine (**Supplementary Fig. 8A–B**, saline: *r*(12) = -0.14, *p* = 0.63, morphine: *r*(8) = 0.075, *p* = 0.84). Taken together, these results indicate that contextual conditioning with opioids leads to the formation and expression of reward memory, as well as the emergence of risk-taking behavior during approach-avoidance conflict.

### Prelimbic cortex neurons show suppressed firing rates following acute, but not repeated, morphine administration

To determine if opioid-induced risk-taking behavior is associated with changes in PL neuronal activity after opioid administration, we used *in vivo* single-unit electrophysiology to record PL neurons at different time points throughout the course of our behavioral experiments. Our first question was whether acute administration of morphine affects PL neuron firing rates in freely-moving rats. To answer this question, we performed recordings before and after the first morphine injection on the first day of conditioning (**Fig. 3A *left***). We used a Z-score-based method of classifying neurons as responsive given their firing rate changes pre- to post-injection (Z-score < -1.96 for inhibition, *p* < 0.05; or Z-score > 2.58 for excitation, *p* < 0.01). Morphine acutely suppressed PL neuron firing rates in both Risk-Avoiders and Risk-Takers (13% or 15% of the neurons, respectively), and these proportions of inhibited cells were significantly greater than the proportion of cells that were inhibited in saline-treated rats (5.5%; **Fig. 3B and 3D–E**, Fisher’s Exact test, saline vs. Risk-Takers: *p* = 0.009). In addition, Risk-Avoiders showed a proportion of PL neurons that responded with increased firing rates to acute morphine injection (**Fig. 3B and 3D–E**, Fisher’s Exact test, saline vs. Risk-Avoiders: *p* = 0.002), an effect that was not observed in Risk-Takers (**Fig. 3B and 3D–E**, Fisher’s Exact test, saline vs. Risk-Takers: *p* = 0.31).

**Figure 3.**
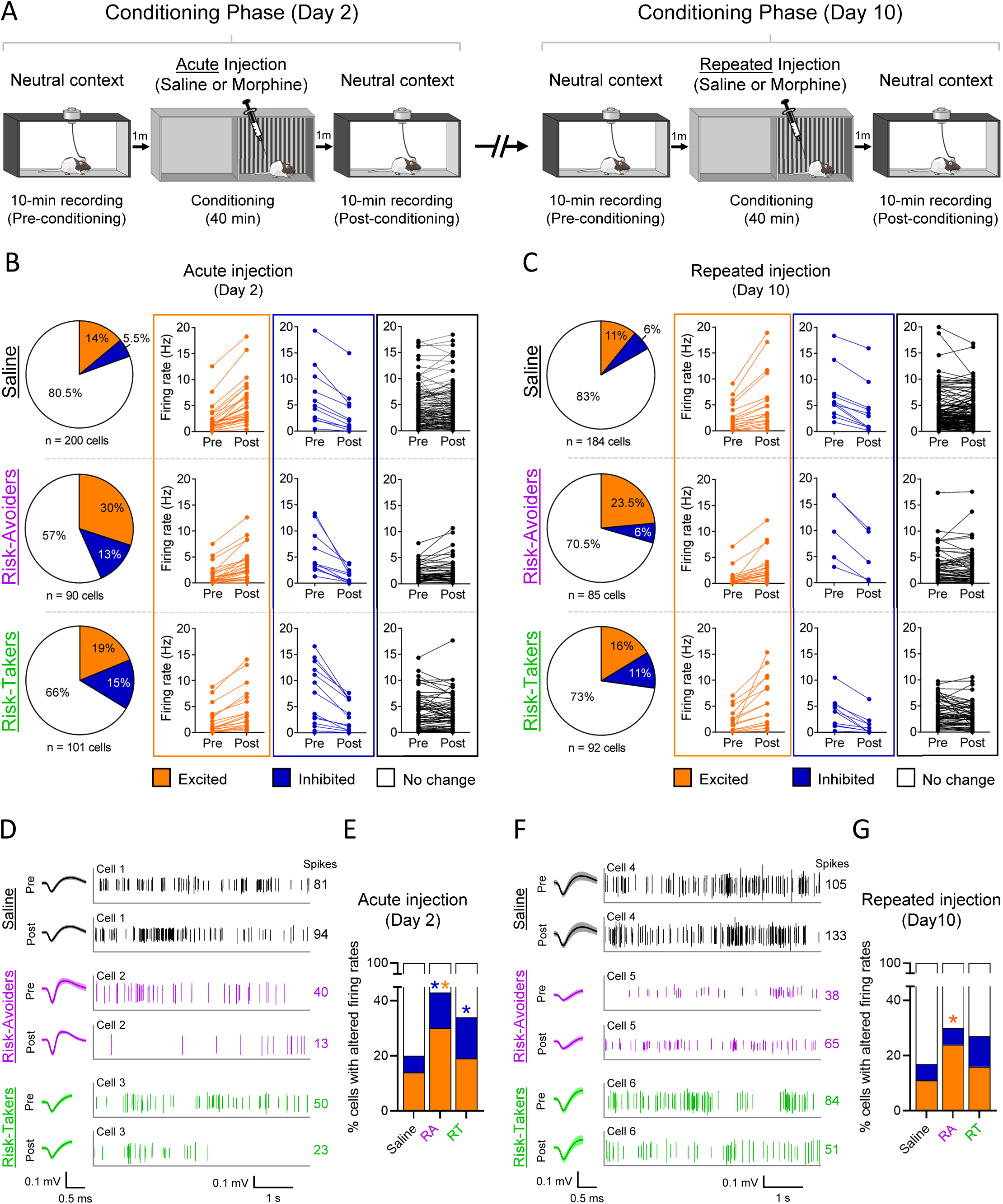
Morphine-induced PL inhibition is lost after conditioning in both Risk-Avoiders and Risk-Takers, but persistent PL excitation is exclusive to Risk-Avoiders. **A)** Timeline of recordings of spontaneous single-unit activity in PL after acute (Day 2) and repeated (Day 10) administration of saline or morphine (10 mg/kg, subcutaneous). **B)** Representations of cells excited (orange), inhibited (dark blue), or exhibiting no change (black/white) in response to acute drug administration (Z-scores used for response classification [excited, >2.58; inhibited, <-1.96]; cells with firing rates >20 Hz not shown [Saline = 7, Risk-Avoiders = 3, Risk-Takers = 3]). As compared to saline administration, acute morphine administration resulted in a greater number of cells showing increased firing rates in Risk-Avoiders (Fisher’s Exact test, ratio of cells excited to not excited, *p* = 0.002), and decreased firing rates in both Risk-Avoiders and Risk-Takers (Fisher’s Exact tests, ratio of cells inhibited to not inhibited; Risk-Avoiders: *p* = 0.0327; Risk-Takers: *p* = 0.0089). **C)** Representations of cells excited (orange), inhibited (dark blue), or exhibiting no change (black/white) in response to acute drug administration (Z-scores used for response classification [excited, 2.58; inhibited, -1.96]; cells with firing rates >20 Hz not shown [Saline = 3, Risk-Takers = 1]). On the final drug administration day, morphine failed to suppress PL cell firing rates beyond what was observed after saline administration (Fisher’s Exact tests, ratio of cells inhibited to not inhibited; Risk-Avoiders: *p* = 1.000; Risk-Takers: *p* = 0.156). However, in Risk-Avoiders, increased PL cell firing rates in response morphine administration were maintained relative to animals that were administered saline (Fisher’s Exact test, ratio of cells excited to not excited, *p* = 0.0091) **D)** Example waveforms and spike raster plots (5s samples from 300s to 305s during recordings) of two representative cells, one from either group, during baseline and after saline or morphine administration on Day 2 (no change cell from Saline group; inhibited cells from Risk-Avoider and Risk-Taker groups). Numbers to the right of spike raster plots denote the quantity of spikes shown. **E)** Relative percentages of cells on Day 2 that were inhibited (Fisher’s Exact tests, RAs vs. saline: *p* = 0.033, RTs vs. saline: *p* = 0.009, RAs vs. RTs: p = 0.84), excited (Fisher’s Exact test, RAs vs. saline: *p* = 0.002, RTs vs. saline: *p* = 0.31, RAs vs RTs: *p* = 0.09), or exhibited no change in response to saline or morphine administration (\**p* < 0.05). **F)** Example waveforms and spike raster plots (5s samples from 300s to 305s during recordings) of two representative cells, one from either group, during baseline and after saline or morphine administration on Day 10 (no change cells from Saline and Risk-Taker groups; excited cell from Risk-Avoider group). Numbers to the right of spike raster plots denote the quantity of spikes shown. **G)** Relative percentages of cells on Day 10 that were excited, inhibited, or exhibited no change in response to saline or morphine administration. Injections failed to result in significant inhibition (Fisher’s Exact tests, RAs vs. saline: *p* = 1.0, RTs vs. saline: *p* = 0.16, RAs vs. RTs: *p* = 0.29), but continued to result in significant excitation solely in the Risk-Avoider group (Fisher’s Exact tests, RAs vs. saline: *p* = 0.009, RTs vs. saline: *p* = 0.25, \**p* = 0.0091).

To determine if morphine’s effects on PL firing rates persist across conditioning sessions, we also performed recordings before and after the final administration of morphine on the last day of drug conditioning (**Fig. 3A *right***). After the final injection, morphine failed to suppress PL firing rates in comparison to repeated saline administration (**Fig. 3C and 3F–G**, Fisher’s Exact tests, saline vs. Risk-Avoiders: *p* = 1.0, saline vs. Risk-Takers: *p* = 0.16). Interestingly, increases in neuronal firing rates observed in Risk-Avoiders during acute morphine administration were maintained during the final morphine conditioning day, differing again from Risk-Takers, which did not show increased neuronal firing (**Fig. 3C and 3F–G**, Fisher’s Exact tests, saline vs. Risk-Avoiders: *p* = 0.009, saline vs. Risk-Takers: *p* = 0.25). These results suggest that although morphine initially suppresses PL activity, there is adaptation to the repeated effects of morphine that renders PL neurons insensitive to this firing rate suppression by the drug. Furthermore, the persistent increase in PL firing rates in Risk-Avoiders following morphine administration during conditioning may underlie the failure of these rats to both form strong contextual drug-reward memories and, ultimately, to exhibit risk-taking behavior during the conflict test.

### The PL neurons of Risk-Takers exhibit enhanced spatial representation of the drug-paired side during conflict

Drug-reward memories are strongly tied to the environmental context in which the drug is used, and re-exposure to this context is a robust driver of drug-seeking behavior (Wikler, 1973; O’brien et al., 1992; Crombag and Shaham, 2002; Hyman, 2005; Crombag et al., 2008; Perry et al., 2014). Information regarding the spatial location of a drug-associated context is represented as spatial maps in the brain, particularly through activity of populations of neurons in the hippocampus (Trouche et al., 2016; Xia et al., 2017; Sjulson et al., 2018; Sun and Giocomo, 2022). Spatial information is also represented by neurons in the prefrontal cortex, where this representation is believed to be important for goal-directed decision making (Poucet and Hok, 2017; Ito et al., 2018; Patai and Spiers, 2021). While PL neurons have been shown to encode the environmental location of natural rewards (Poucet, 1997; Poucet et al., 2004; Hok et al., 2005; Powell and Redish, 2014; Murugan et al., 2017; Hunsaker and Kesner, 2018; Hasz and Redish, 2020a, 2020b; Diehl and Redish, 2023), it is unknown how contextual drug memories are processed in this region.

To define patterns of spatial representation of opioid context in PL after conditioning, we used a behavioral pose estimation algorithm (DeepLabCut; Mathis et al., 2018) in conjunction with single-unit recordings to align PL neuronal firing rates with rats’ head positions in the apparatus during the preference and conflict tests. (**Fig. 4A**) Our analysis revealed that PL neurons exhibited location-specific activity, with distinct subsets of neurons showing either inhibitory or excitatory responses when the rats occupied specific areas of the apparatus (**Fig. 4B**, Z-score < -1.96 for inhibition, *p* < 0.05; or Z-score > 2.58 for excitation, *p* < 0.01). Notably, Risk-Takers demonstrated a higher percentage of neurons that responded to the paired side during the conflict test compared to both Risk-Avoiders and saline controls (**Fig. 4C**, Chi-square test, *p* = 0.035). Furthermore, Risk-Takers exhibited a significant increase in the number of neurons responsive to the paired side from the preference test to the conflict test (**Fig. 4C**, Fisher’s Exact test, ratio of responsive cells to non-responsive cells, preference test vs. conflict test, *p* = 0.021), which may have been driven by inhibited neurons as the proportion of excited neurons did not change between the two test phases in this group (**Fig. 4D–F**, Fisher’s Exact test, ratio of excited cells to non-excited cells, preference test vs. conflict test, *p* = 0.103). This increase in inhibited PL cells in Risk-Takers is especially interesting when compared to the saline group, which exhibited a lower proportion of inhibited PL cells during the conflict test compared to the morphine-treated groups (**Fig. 4D**, Chi-square test, *p* = 0.036). These findings suggest that Risk-Takers have enhanced neural representation of the drug-associated context in PL during motivational conflict, which is driven by inhibitory signaling and may underlie the persistent risk-taking behavior observed in this group.

**Figure 4.**
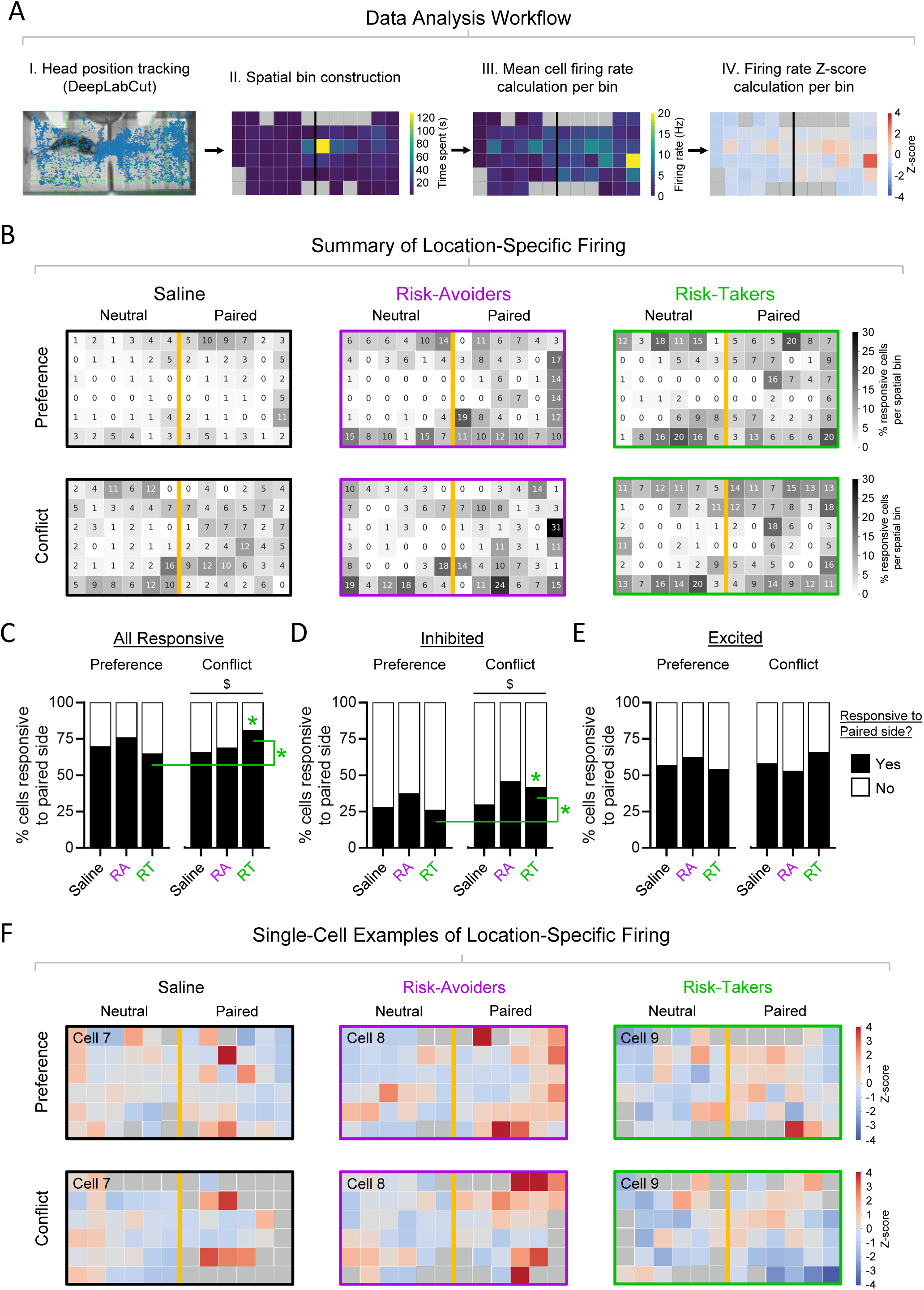
The PL neurons of Risk-Takers exhibit enhanced spatial representation of the drug-paired side during conflict. **A)** Schematics summarizing the data analysis pipeline used to determine spatially-defined firing of PL neurons in the CPP apparatus. I) Head positions of rats in the apparatus during the preference and conflict tests were tracked using DeepLabCut pose estimation software. II) For each test video, the apparatus was segmented into 72 spatial bins and the amount of time each animal spent in each bin was calculated (gray bins are those which the animal did not enter during the test). III) The mean firing rate for each PL neuron was calculated when the animal’s head was located in each spatial bin (gray bins are those which the animal was present for less than 5 video frames). IV) The firing rates for each neuron in each spatial bin were normalized to the average firing rate of the same cell during the entire session to determine response classification (excited, Z-score >2.58, p<0.01; inhibited, Z-score < -1.96, p<0.05). **B**) Plots showing the percentages of PL cells within each group that responded with significant firing rate changes in specific spatial bins of the apparatus during the preference and conflict tests. Neurons responding in more than one spatial bin are included in the quantification. **C**) Percentages of PL cells that exhibited significant firing rate changes (inhibited or excited) in any spatial bin within the paired side of the apparatus during the preference or conflict tests. Risk-Takers exhibited an increase in the percentage of paired side-responsive PL cells during the conflict test (Fisher’s Exact tests, ratio of responsive cells to non-responsive cells, preference test vs. conflict test, Saline: *p* = 0.552, RA: *p* = 0.454, RT: *p* = 0.021), and this percentage was greater than either Saline or Risk-Avoider rats (Chi-square test, ^$^*p* = 0.035). **D**) Percentages of PL cells that significantly decreased their firing rates in any spatial bin within the paired side of the apparatus during the preference or conflict tests. Risk-Takers exhibited an increase in the percentage of paired side-inhibited PL cells during the conflict test (Fisher’s Exact tests, ratio of inhibited cells to non-inhibited cells, preference test vs. conflict test, Saline: *p* = 0.807, RA: *p* = 0.398, RT: *p* = 0.032). Saline-treated rats displayed a smaller proportion of paired side-inhibited PL cells during the conflict test than either Risk-Avoiders or Risk-Takers (Chi-square test, preference test: *p* = 0.253, conflict test: ^$^*p* = 0.036). **E**) Percentages of PL cells that significantly increased their firing rates in any spatial bin within the paired side of the apparatus during the preference or conflict tests. None of the three groups showed changes in the proportion of paired side-excited PL cells between the preference and conflict tests (Fisher’s Exact tests, ratio of excited cells to non-excited cells, preference test vs. conflict test, Saline: *p* = 0.911, RA: *p* = 0.312, RT: *p* = 0.103), nor changes between groups during the preference or conflict tests (Chi-square tests, preference test: *p* = 0.518, conflict test: *p* = 0.191). **F**) Single-cell examples from each of the three groups showing spatially-constrained firing properties during the preference test (*top row*) and conflict test (*bottom row;* gray bins are those which the animal was present for less than 5 video frames).

### PL neurons of Risk-Takers show persistent inhibitory responses upon initiating exploration of the drug-paired side during conflict

In a CPP task, rats must rely on contextual information to identify a drug-associated context while freely exploring the apparatus (Green and Bardo, 2020). PL neurons can encode reward locations in an environment, and rapidly adapt their firing rates when reward locations change and animals adjust goal-directed navigational strategies to obtain these rewards (Poucet, 1997; Hok et al., 2005; Erdem and Hasselmo, 2012; Powell and Redish, 2014, 2016; Ito et al., 2015, 2018; Murugan et al., 2017; Hasz and Redish, 2020a, 2020b; Diehl and Redish, 2023). To determine if individual PL neurons display firing rate changes upon exploration of the drug-associated context, we aligned neuronal spiking activity to the moment at which rats’ heads entered the drug-paired side of the apparatus (**Fig. 5A**). We then applied spectral clustering, an unsupervised learning algorithm, based on the normalized firing rates of the entire session to identify groups of PL neurons with similar temporal firing patterns during paired-side entries in both preference and conflict tests (319 neurons from 22 rats). From this analysis emerged 7 distinct clusters of PL neurons, each with unique response profiles (**Fig. 5B–D**). We confirmed the segregation of clusters using dimensionality reduction (tSNE transformation) to visualize the data in a 2D plot and compare tSNE scores across each cluster pair (**Supplementary Fig. 9A–B**). We found that Cluster 2, which included neurons with higher firing rates during paired-side entries in the preference test vs. the conflict test, represented a greater ratio of neurons recorded from morphine-treated rats (Risk-Avoiders and Risk-Takers) than from saline-treated rats (**Fig. 5B**, Fisher’s Exact test, saline vs. morphine: *p* = 0.038). This result suggests a dynamic processing of drug-associated context in PL that reflects the transition from drug seeking to approach-avoidance conflict.

**Figure 5.**
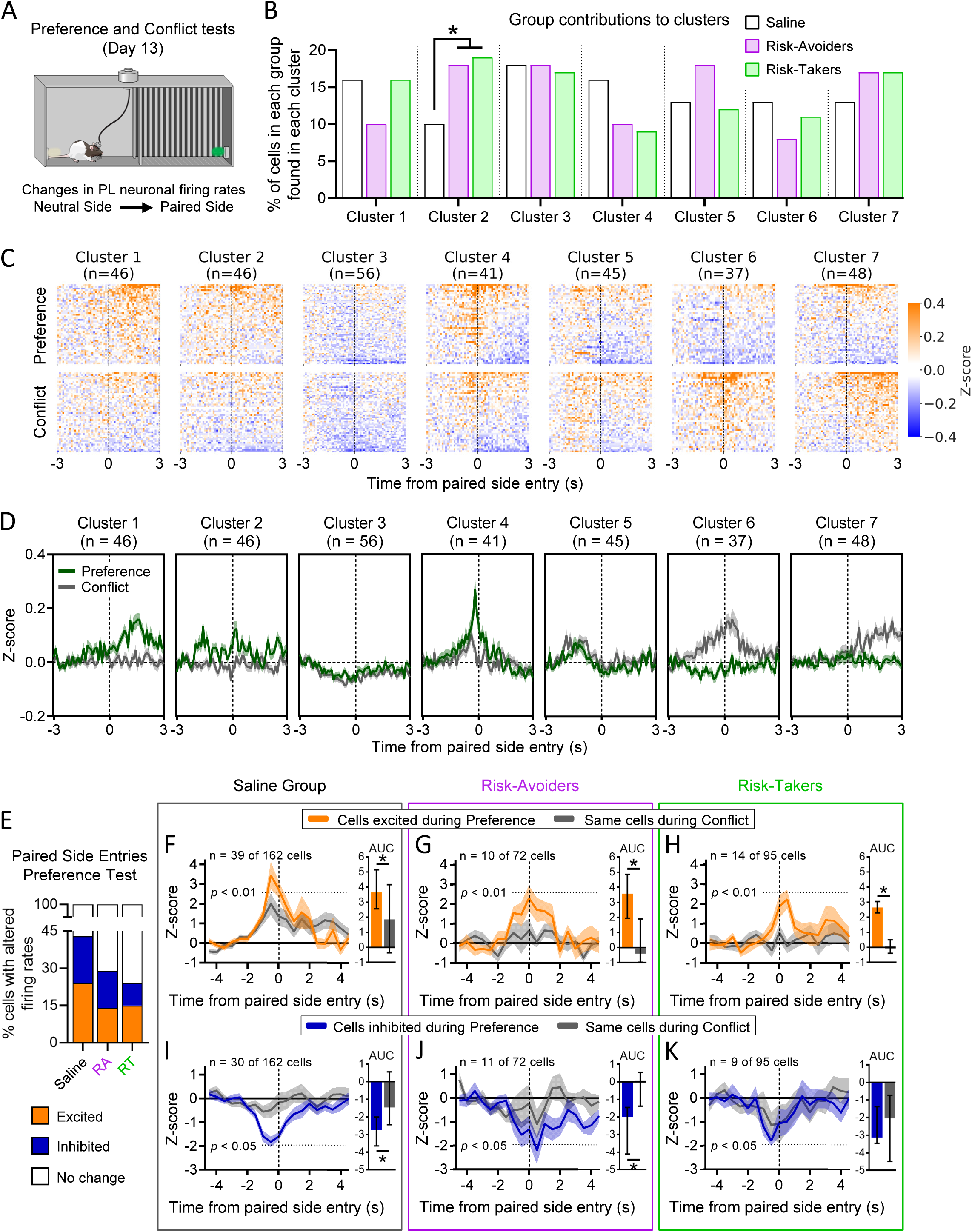
PL neurons of Risk-Takers show persistent inhibitory responses upon initiating exploration of the drug-paired side during conflict. **A**) Schematic diagram showing the behavior (paired-side head entries) to which neuronal activity was aligned for the following analyses. **B**) Percentages of cells identified from recordings from each group that were included in Clusters 1 through 7 after spectral clustering. A larger proportion of PL cells in morphine-treated rats (Risk-Avoiders and Risk-Takers) exhibited paired side entry responses consistent with Cluster 2 compared to Saline-treated rats (Fisher’s Exact test, saline vs. morphine: *p* = 0.038). **C**) Single-unit peri-event raster plots and **D**) mean peri-event time histograms showing firing rate changes of PL cells relative to paired-side entries in the preference and conflict tests. All data shown as Z-scores. **E**) Percentages of cells showing excitation (Z-score > 2.58), inhibition (Z-score < - 1.96), or no response to paired side entries during the preference test (saline-treated group, n = 9 rats: 39/162 [24%] cells excited, 30/162 [19%] cells inhibited; RA group, n = 6 rats: 10/72 [14%] cells excited, 11/72 [15%] cells inhibited; RT group, n = 7 rats: 14/95 [15%] cells excited, 9/95 [9%] cells inhibited; Fisher’s Exact tests, RAs vs. saline: *p* = 0.059, RTs vs. saline: *p* = 0.0031, RAs vs. RTs: *p* = 0.48). **F–H**) Graphs representing firing rate changes of PL cells in Saline (F; n = 9 rats), Risk-Avoider (G; n = 6 rats), and Risk-Taker (H; n = 7 rats) groups that showed excitatory spontaneous activity (Z-score > 2.58) when animals crossed into the paired side during the preference test (orange) compared to firing rates of the same cells when animals crossed into the paired side during the conflict test (charcoal). Inset bar graphs show differences in the total areas under the curves between test stages 500 msec before and after line crossings (*\*p* < 0.05). PL neurons that responded to paired side entries during the preference test with increased firing rates did not respond to paired side entries during the conflict test in Saline-treated rats (AUC: Shapiro-Wilk normality test, *p* < 0.0001; Wilcoxon test, *p* = 0.013), Risk-Avoiders (AUC: Shapiro-Wilk normality test, *p* < 0.01; Wilcoxon test, *p* = 0.027) or Risk-Takers (AUC: paired Student’s t-test, *p* = 0.0013). **I–K**) Graphs representing firing rate changes of PL cells in Saline (I; n = 9 rats), Risk-Avoider (J; n = 6 rats, AUC), and Risk-Taker (K; n = 7 rats) groups that showed inhibitory spontaneous activity (Z-score < -1.96) when animals crossed into the paired side during the preference test (dark blue) compared to firing rates of the same cells when animals crossed into the paired side during the conflict test (charcoal). Inset bar graphs show differences in the total areas under the curves between test stages 500 msec before and after line crossings (*\*p* < 0.05). PL neurons that responded to paired side entries during the preference test with decreased firing rates did not respond to paired side entries during the conflict test in either Saline-treated rats (AUC: Shapiro-Wilk normality test, *p* < 0.01; Wilcoxon test, *p* = 0.0002) or Risk-Avoiders (AUC: Shapiro-Wilk normality test, *p* < 0.01; Wilcoxon test, *p* = 0.032). However, in Risk-Takers, inhibited paired side entry-responsive PL cells showed similar spatial firing rate changes during both the preference and conflict tests (AUC: Shapiro-Wilk normality test, *p* < 0.01; Wilcoxon test, *p* = 0.73). Data are shown as median ± interquartile range for Wilcoxon tests (F–G, I–K) and mean ± SEM for paired Student’s t-test (H).

To examine possible conflict-associated shifts in contextual processing by PL neurons in greater detail, we visually categorized neurons based on their individual firing rate responses to paired-side head entries. In all three groups of rats, we identified distinct populations of PL neurons that were either excited (Z-score > 2.58, *p* < 0.01) or inhibited (Z-score < -1.96, *p* < 0.05) during paired-side head entries in the preference test, with lower proportions of responsive neurons in Risk-Takers than in saline-treated rats (**Fig. 5E**, Fisher’s Exact test, saline vs. Risk-Takers: *p* = 0.0031), suggesting that individual PL neurons display firing rate changes upon exploration of both drug-associated and neutral contexts.

After we identified PL neurons that exhibited firing rate changes associated with paired-side head entries, we investigated whether these signals of exploration were altered in the presence of cat odor when rats adopted new behavioral strategies (*i.e*., risk avoiding or risk taking). To answer this question, we compared the activity of paired side-responsive neurons during the preference test, described above, to their activity when aligned to paired-side entries during the conflict test (**Fig. 5F–K**). In saline-treated rats, PL neurons that were excited (**Fig. 5F**, **Supplementary Fig. 10A**) or inhibited (**Fig. 5I**, **Supplementary Fig. 10B**) in response to paired-side entries during the preference test showed attenuated responses (*i.e.*, less excitation or inhibition) to paired-side entries during the conflict test (excited: area under the curve [AUC], Wilcoxon test, *p* = 0.013; inhibited: AUC, Wilcoxon test, *p* = 0.0002). This finding suggests that in PL, neuronal signals of exploration of a neutral context are suppressed once the context becomes associated with an aversive stimulus.

We next performed the same analyses in Risk-Avoiders and Risk-Takers to determine if the motivational conflict induced by the presence of cat odor would modify PL neuronal activity during the exploration of the drug-associated context. As in saline-treated rats, excitatory paired-side entry responses were suppressed during the conflict test in both Risk-Avoiders (**Fig. 5G**, AUC, Wilcoxon test, *p* = 0.027; **Supplementary Fig. 10C**) and Risk-Takers (**Fig. 5H**, AUC, paired Student’s t-test, *p* = 0.0013; **Supplementary Fig. 10E**), suggesting that in PL, excitatory signals of morphine-associated context exploration are lost when the context acquires conflicting valences. Because inhibitory PL responses to paired-side exploration were attenuated during the conflict test in saline-treated rats (**Fig. 5I**, described in the previous paragraph), we asked if the same would be true for morphine-treated rats. In Risk-Avoiders, inhibitory responses to paired-side exploration were attenuated during the conflict test in a manner similar to that observed in saline-treated rats (**Fig. 5J**, AUC, Wilcoxon test, *p* = 0.032; **Supplementary Fig. 10D**). In contrast, Risk-Takers differed from both Risk-Avoiders and saline-treated rats in that PL neurons that were inhibited during paired side exploration in the preference test showed similar inhibitory responses during the conflict test (**Fig. 5K**, AUC, Wilcoxon test, *p* = 0.73; **Supplementary Fig. 10F**). This persistent inhibitory signaling of the drug-associated context from the preference to the conflict tests in Risk-Takers mirrors the persistent exploration of the drug-paired side observed during the conflict test in this subgroup of rats. Taken together, our results indicate that persistent inhibitory signaling of the drug-associated context in a subset of PL neurons during conflict may underlie increased risk taking following opioid exposure.

## DISCUSSION

We established a novel CPP procedure to study drug-induced risk-taking behavior in rats. We found that conditioning with opioid drugs led to context-dependent risk-taking behavior characterized by persistent CPP, despite the coincident presence of predator threat, in one subset of rats (Risk-Takers), but not in another (Risk-Avoiders). Through *in vivo* single-unit recordings, we observed that a significant proportion of PL neurons exhibited suppressed firing rates in response to acute morphine injection, but not following repeated morphine exposure. We also observed that Risk-Avoiders, exclusively, exhibited increased PL firing rates upon morphine exposure, a response that persisted throughout conditioning. Further analysis revealed distinct populations of PL neurons that showed firing rate changes consistent with representation of environmental cues relative to the rat’s location within the apparatus, with a greater number of neurons encoding the drug-paired context in Risk-Takers than in Risk-Avoiders or saline control rats during conflict. Additionally, we found subsets of PL neurons that displayed firing rate changes associated with rats’ transitions between contexts within the apparatus. In both Risk-Avoider and saline groups, PL neurons that showed suppressed firing rates when rats entered the drug-paired context during the preference test lost these inhibitory responses during the conflict phase in the presence of predator odor. However, this inhibitory signal of drug-paired side exploration persisted in Risk-Takers during the conflict test, mirroring the risk-taking behavior observed in this group of rats. Together, our results suggest a mechanism of opioid-induced risk-taking behavior that involves failed prefrontal signaling of threat when drug- and threat-associated cues occur in the same context.

### Individual phenotypes and sex differences in opioid-induced risk-taking behavior

In our experiments, conditioning with opioid drugs (morphine or fentanyl) led to the formation of contextual reward memory and subsequent risk-taking behavior. These results resemble those from prior rodent studies using opioid self-administration punished by electrical footshocks, which reported near complete suppression of opioid seeking as a function of footshock intensity (Panlilio et al., 2003, 2005; Peck et al., 2013, 2015; Borges et al., 2022, 2023; Honeycutt et al., 2022; Mathieson et al., 2022; Fredriksson et al., 2023). However, other studies have reported incomplete suppression of opioid seeking or divergent behavioral responses that resulted in subgroups of either punishment-sensitive or punishment-resistant rats (Porter-Stransky et al., 2017; Blackwood et al., 2020). Notably, our model differs from footshock studies in that we used a non-nociceptive aversive stimulus (cat saliva) to avoid potential confounds associated with opioids’ documented effects on nociceptive sensitivity (Liebmann et al., 1997; Lee et al., 2011). Other rodent studies have attempted to model risky opioid seeking using non-nociceptive stimuli by employing air puffs (Skupio et al., 2017), the bitterant quinine during oral self-administration (Monroe and Radke, 2021), or co-administration of histamine (Minervini et al., 2021; Honeycutt et al., 2022) as response-contingent aversive stimuli. While these studies did not explore individual behavioral differences, none of the non-nociceptive stimuli employed were sufficient to eliminate opioid seeking overall. Whether these non-nociceptive stimuli can completely suppress opioid seeking if administered with sufficient intensity remains unknown.

In our study, we also identified sex differences in risk-taking behavior following comparable doses of morphine. Particularly, conditioning with 10 mg/kg of morphine led to CPP and risk-taking behavior during conflict in male but not female rats. Our CPP results are consistent with some previous studies (Wang et al., 2019; Gaulden et al., 2021), but not others which have reported either similar levels of opioid CPP in male and female rodents (Randall et al., 1998; Campbell et al., 2000; Randesi et al., 2019; Barattini et al., 2023), or greater sensitivity to the rewarding properties of opioids in females (Cicero et al., 2000; Karami and Zarrindast, 2008; Timár et al., 2010; Roversi et al., 2016; Ryan et al., 2018; Bellamy et al., 2019; Reich et al., 2019). While the cause of these discrepancies is unclear, differences across these studies in factors such as opioid type/dose, rodent strain, conditioning protocol, and estrous cycle phase in females may explain the inconsistent results. In contrast to morphine, we did not observe sex differences in CPP or risk-taking behavior following fentanyl conditioning. Given that sex differences in fentanyl reward appear to be dose dependent (Gaulden et al., 2021; Barattini et al., 2023; Little and Kosten, 2023), it is possible that a higher dose of morphine than the one we used in our study may elicit risk taking in female rats. Thus, the interplay among opioid reward, context, and risk taking is a multifaceted factor, and our results underscore the importance of sex as a critical variable in understanding these interactions.

### Changes in PL neuronal activity in response to repeated opioid administration

We observed a reduction of PL firing rates following acute morphine administration, consistent with previous findings showing that opioids suppress neuronal activity in mPFC (Giacchino and Henriksen, 1996). However, following repeated morphine injections during CPP, PL neurons underwent adaptation and no longer exhibited suppressed firing rates in response to opioid administration. Prior studies measuring the firing activity of PL neurons in response to acute opioid administration have resulted in contrasting findings; while some studies have reported increased excitability (Sun et al., 2011; Sun and Laviolette, 2012; Jiang et al., 2021), others have shown inhibitory effects (Lee et al., 1999). In comparison, studies examining neural adaptations associated with repeated opioid exposure have revealed complex changes in PL neurons, including alterations in synaptic strength, dendritic spine morphology, glutamate receptor expression, among others (Ballesteros-Yáñez et al., 2007; Shen and Kalivas, 2013; Zhu et al., 2020; Anderson et al., 2021; Kokane et al., 2023; Siemsen et al., 2023). Our additional result that morphine-induced PL excitation occurs exclusively in Risk-Avoiders, without diminishing with repeated administration, suggests that both initial and long-term neuronal responses to opioids in PL are important for predicting subsequent risk-taking behavior (Blackwood et al., 2020). Together, our findings demonstrate that opioid-induced plasticity in PL is associated with individual differences in risk-related decision-making during approach-avoidance conflict.

### Encoding of drug-associated spatial information in PL neurons

Combining single-unit recordings from PL neurons with behavioral pose estimation using DeepLabCut (Mathis et al., 2018) allowed us to align neuronal firing patterns with the spatial location of the rats during our tests. Our observation that PL neurons exhibit location-specific activity, with distinct subsets of neurons increasing or decreasing their firing rates when rats occupied specific subregions within the apparatus, is consistent with previous studies showing that PL neurons encode spatial information, particularly in relation to the location of natural rewards such as food (Poucet, 1997; Hok et al., 2005; Powell and Redish, 2014; Murugan et al., 2017; Hasz and Redish, 2020a, 2020b; Diehl and Redish, 2023).

Our study also demonstrates that PL neurons process spatial information in the context of drug reward, as the proportion of cells showing location-specific firing in the drug-paired side increases from the preference to the conflict test in Risk-Takers. This increase was driven by an elevated number of inhibited neurons during the conflict test, a phenomenon not observed in Risk-Avoiders or saline-treated controls. These results suggest that Risk-Takers may require a heightened level of neural processing by the PL to resolve a motivational conflict. Notably, a recent study has reported the appearance of hippocampal place cells encoding location within the drug-paired side of a CPP apparatus following morphine conditioning in mice, which the authors attributed to a learned over-representation of reward location (Sun and Giocomo, 2022). In our study, the increased number of paired side-responsive PL cells in Risk-Takers may reflect an enhanced neural representation of the drug-associated context during motivational conflict, which could underlie their persistent risk-taking behavior even in the face of potential threats.

### PL neuronal activity during opioid-induced risk-taking behavior

Using single-unit recordings from PL neurons during our conflict test, we found an inhibitory neuronal signal of drug context that persisted during conflict in Risk-Takers, but was abolished during conflict in Risk-Avoiders. Reduced PL activity has been previously reported in risk-taking rats during a sucrose-approach versus threat-avoidance conflict test (Bravo-Rivera et al., 2021). Recent work from our lab identified a similar pattern of inhibition in PL glutamatergic neurons of risk-taking rats during an approach-avoidance conflict test, and optogenetic-mediated inhibition of these cells during food cue presentations in risk-avoiding rats restored reward seeking despite threat (Fernandez-Leon et al., 2021). Further support for a role of PL in risky drug seeking comes from a separate study showing reduced activity of the PL to ventral striatum pathway in footshock-resistant rats during methamphetamine seeking (Hu et al., 2019), which is consistent with earlier reports of PL hypofunction in cocaine seeking despite punishment (Chen et al., 2013; Limpens et al., 2015; Smith and Laiks, 2018). While we observed a subpopulation of PL neurons with persistent inhibition during risk taking in our model, we also found neurons with concurrent excitatory responses, as well as a nonresponsive majority (see **Fig. 5E**).

Given the intricacy of PL dynamics, the unknown genetic identity of the neurons involved, and our model’s focus on exploratory behavior rather than temporally pre-defined cue responses, we opted to forego chemogenetic or optogenetic manipulations that might bias PL activity across temporal and spatial scales incongruent with the single-unit activity we observed. Future studies in our lab aim to adapt our cat odor conflict model to facilitate cued drug seeking alongside appropriate neural manipulations for causal testing. These studies will add to the growing literature that suggests the neurophysiological underpinnings of suppressed drug seeking differ depending on whether the suppression is voluntary or not (Fredriksson et al., 2021b, 2023; Negishi et al., 2024). In addition to PL’s role in models of risky drug seeking, there may also be value in investigating the contribution of PL in other opioid-sensitive models of risk taking, especially those that involve seeking natural rewards (Vollmer et al., 2022; Wheeler et al., 2023).

Collectively, our findings establish a prefrontal neural signal of risky opioid seeking in a threatening context. Our results can inform translational studies aimed at identifying prefrontal neural biomarkers in humans who might be prone to risky opioid use, as well as clinical studies targeting prefrontal activity as a therapeutic approach for people diagnosed with opioid use disorder.

## ACKNOWLEDGEMENTS

We would like to thank our colleagues who provided valuable feedback on earlier versions of this manuscript, including Christian Bravo-Rivera, Freddyson Martinez-Rivera, James Otis, and Leandro Vendruscolo. We thank Nikita Watson and Sharon Gordon for their technical and administrative assistance.

## AUTHOR CONTRIBUTIONS

C.Q., A.M.V., and D.S.E. performed the behavioral experiments. C.Q. and G.A.-M. performed the stereotaxic surgeries. C.Q. and A.M.V. performed the single-unit recording experiments. C.Q., M.M., E.P.B, C.L.C., X.O.Z., and F.H.D.-M. analyzed the single-unit recording experiments. M.M. performed the behavioral analyses using DeepLabCut and developed the data analysis pipeline for spatial representation. All the authors participated in the design of the experimental protocols and interpretation of the data. C.Q. and F.H.D.-M. prepared the manuscript with comments from all the co-authors.

## FUNDING

This study was supported by a National Institutes of Health (NIH) grant F31-DA-055469 to C.Q., the Russell & Diana Hawkins Family Foundation Discovery Fellowship and the Dr. John J. Kopchick Fellowship to X.O.Z., and an NIH grant R00-MH105549, an NIH grant R01-MH120136, an NIH grant R01-MH131570, a Brain & Behavior Research Foundation grant (NARSAD Young Investigator), and a Rising STARs Award from UT System to F.H.D.-M.

## COMPETING INTERESTS

The authors declare no competing interests.

## SUPPLEMENTARY METHODS SECTION

### Cat odor collection and preparation

Cat saliva was collected from anesthetized cats at a local veterinary clinic (https://www.snapus.org) as described in our previously published work (Engelke et al., 2021). Briefly, a filter paper strip was placed under the tongue of an anesthetized cat for approximately 10 minutes. These saliva-soaked strips were then weighed, transported to the lab at 4°C, and stored at -20°C until used in experiments. The collected saliva represented diverse cat demographics, including different sexes, strains, and ages, with prior studies suggesting consistent defensive response patterns across these variables (Muñoz-Abellán et al., 2010).

### Behavioral tasks

#### Conditioned place preference and conflict tests

Rats were exposed to a two-chamber apparatus (31 cm length X 31 cm width X 40 cm height, each side) with each chamber having distinct visual (solid vs. striped walls) and tactile (drilled vs. smooth floors) cues. The two chambers were connected by a corridor that could accommodate a removable barrier. Throughout the experiment, a clear plexiglass lid was placed over both chambers to prevent climbing or escape. On Day 1, saline-saturated filter paper strips were positioned in metal mesh housings (5 cm diameter) and affixed to the far walls of both sides of the apparatus in preparation for the conflict test. Rats were allowed to freely explore both sides of the apparatus for 10 min and time spent on either side was recorded as a baseline measure of side preference. Baseline preference was defined as greater than 55% of time spent in one side. On Day 2, both the filter paper strips and the metal mesh frames were removed from the apparatus and the conditioning phase began. During conditioning, rats received subcutaneous injections of either morphine (5 or 10 mg/kg, 0.5 or 1 ml/kg), fentanyl (0.02 or 0.04 mg/kg, 0.4 or 0.8 ml/kg), or saline (1 ml/kg) and were confined for 40 min to the side of the apparatus preferred least at baseline (i.e., a biased design). These doses were chosen because they were previously reported in the literature to produce CPP in rats (Mucha and Herz, 1985; Finlay et al., 1988; Bardo et al., 1995; Miller and Nation, 1997; Vitale et al., 2003; Milekic et al., 2006; Martínez-Rivera et al., 2016). On Day 3, rats were subcutaneously injected with saline and confined to the opposite side of the apparatus for 40 min. Drug and saline conditioning continued every other day for 10 days (Days 2-11) for a total of 5 drug and 5 saline pairings. 48 hours after the final conditioning day, filter paper strips saturated with saline were affixed to the far walls of both sides of the apparatus (as on Day 1) and rats were allowed to freely explore both sides of the apparatus for 10 min (Day 13 – Preference Test). Total time spent in each side was recorded to calculate place preference. Immediately after the preference test, rats were removed from the apparatus and the filter paper strips saturated with saline located in the side of the apparatus previously paired with drug were replaced with filter paper strips saturated with cat saliva. Rats were then placed back into the apparatus and allowed free access to both sides for 10 min (Day 13 – Conflict Test), with time spent in each side being recorded as a measure of preference/aversion.

#### Homecage opioid administration test

On Day 1, rats were exposed to a two-chamber apparatus (described above) and allowed to freely explore both sides for 10 min. Filter paper strips saturated with saline were positioned in metal mesh housings and affixed to the far walls of both sides of the apparatus. Time spent on either side was recorded as a baseline measure of side preference. Baseline preference was defined as greater than 55% of time spent in one side. On Day 2, rats received subcutaneous injections of either morphine (10 mg/kg, 1 ml/kg) or saline (1 ml/kg) and placed back into their homecage. On Day 3, all rats received subcutaneous injections of saline and placed back into their homecage. Drug and saline administration continued every other day for 10 days (Days 2-11) for a total of 5 drug and 5 saline administrations. 48 hours after the final injection day, filter paper strips saturated with saline were positioned in metal meshes and affixed to the far walls of both sides of the apparatus (as on Day 1). Rats were allowed to freely explore both sides of the apparatus for 10 minutes (Day 13 – Pre-Test). Immediately after the pre-test, rats were removed from the apparatus and the filter paper strips saturated with saline located in the side of the apparatus preferred most at baseline were replaced with filter paper strips saturated with cat saliva. Rats were then placed back into the apparatus and allowed free access to both sides for 10 min (Day 13 – Cat Odor Test), with time spent in each side being recorded as a measure of aversion.

#### Stereotaxic surgeries for electrode implantation

Rats were placed under anesthesia using 5% isoflurane by inhalation in an induction chamber. Rats were then placed in a stereotaxic frame (Kopf Instruments) and anesthesia was maintained with 2.5% isoflurane via a nose mask. During the surgery, rats’ respiration and body temperature were closely monitored, and a heating pad was placed under their body. Local anesthesia (bupivacaine, 0.25%, 3 mL) was provided at the incision site. For asepsis, the incision site was sterilized by alternately wiping with iodine and ethanol (70%). Veterinary lubricant ointment was applied to rats’ eyes to prevent excessive dryness during surgeries. An array of 32 microwires (Bio-Signal) was implanted into the right hemisphere aiming at PL by using the following coordinates (relative to the bregma): –2.6 mm anteroposterior (AP); +0.7 mm mediolateral (ML); -3.6 mm dorsoventral (DV). The ground wire was inserted beneath the skull above the left hemisphere and was wrapped around a grounding screw anchored into the skull at the same location. Additionally, six anchoring screws were placed in the skull. The skull, screws, and implanted electrode were coated with C&B Metabond (Parkell) followed by ortho acrylic cement. Two insulated metal hooks were implanted bilaterally into the cement on either side of the array connector to allow firm attachment of the cable during recording. After surgery, the incision site was coated with triple antibiotic ointment and rats were injected with meloxicam (1 mg/kg, subcutaneous) for postoperative analgesia.

### *In vivo* single-unit electrophysiology

#### Neuronal data acquisition and processing

Electrophysiological recordings were obtained from freely-moving rats using a 64-channel neuronal data acquisition system (Omniplex, Plexon). Rat location data obtained from video tracking software (ANY-maze, Stoelting) and single unit recordings were combined in the same file, allowing for correlation of neuronal activity with behavior. The head-mounted array was connected to a 32-channel digital headstage cable (Plexon) and fed into the data acquisition system via a digital headstage processor (Plexon) and a motorized carrousel commutator (Plexon). A band-pass filter (500-5000 Hz) was applied to extracellular recording data. Extracellular waveforms were digitized at 40 kHz and stored locally on a disk. An automated valley-seeking scan algorithm identified spikes, which were then manually sorted using visually represented sort quality metrics (Offline Sorter, Plexon). Single unit identification was based on two-or three-dimensional combinations of three principal components and various waveform features, including peak-valley relationships and spike amplitude. Spontaneous firing rates, drug-evoked responses, and rat location-related responses were calculated using commercially available software (NeuroExplorer, NEXT Technologies).

#### *In vivo* electrophysiological recordings following acute and repeated drug administration

Rats were exposed to a rectangular plexiglass arena (recording arena, 60 cm length X 26 cm width X 40 cm height; Med Associates) while connected to the headstage cable daily for at least 2 days prior to testing to become accustomed to the recording procedure and context. To determine the acute effects of opioid drugs on PL neuronal firing rates, single-unit recordings were performed in the recording arena (separate from the two-chamber conditioned place preference apparatus) on conditioning Day 2 (first drug administration) and Day 10 (fifth and last drug administration). Prior to injection, rats were connected to the headstage cable and placed into the recording arena, after which electrophysiological activity was recorded for 10 min. Rats were then moved to the conditioned place preference apparatus and underwent conditioning (Day 2) for 40min. Immediately after conditioning, rats were reconnected to the headstage cable and placed back into the recording arena, where recording again took place for 10 min. Similarly, rats underwent single-unit recordings to determine the effects of repeated opioid drug administration on PL neuronal firing rates for 10 min prior to and 10 min after the fifth and last drug injection on Day 10 of conditioning. Spontaneous and drug-evoked neural activity were calculated by comparing the frequency of spike trains during the entirety of the pre-test (10 min) to that during the entirety of the post-test (after injection and conditioning; 10 min). Neuronal responses were classified using Z-score values obtained from this comparison. Neurons with firing rate changes eliciting a Z-score >2.58 (*P* < 0.01) were classified as excited, whereas those with firing rate changes eliciting a Z-score <-1.96 (*P* < 0.05) were classified as inhibited.

#### Preference test and conflict test recordings

Head entries were identified via handscoring of videos obtained by video tracking software during recordings (ANY-maze, Stoelting). Because the start times of behavioral videos and electrophysiological recordings were not always identical, temporal alignment for handscoring was determined by calculating differences between rat center point transitions identified by behavioral tracking software and the associated flags that were imbedded in electrophysiological recording files (Omniplex, Plexon). Rat head entry responses were calculated as *Z*-scores normalized to 10 pre-cue bins of 500 ms comprising the -6 s to -1 s period prior to each head entry; to maintain a clean baseline for neuronal activity, head entries that occurred less than 6 s after a previous head entry were excluded from analyses. Neuronal responses were classified using Z-score values obtained from this comparison, considering 2 bins before and 2 bins after each head entry timepoint, using individual time bins of 1 s. Neurons with firing rate changes in any of these bins eliciting a Z-score >2.58 (*p* < 0.01) were classified as excited, whereas those with firing rate changes eliciting a Z-score <-1.96 (*p* < 0.05) were classified as inhibited.

### Histology

Rats were euthanized by intraperitoneal injection of a pentobarbital solution (Fatal-Plus, Vortech Pharmaceuticals) and transcardially perfused with KBPS, followed by perfusion with 10% phosphate buffered formalin. Brains histological processing occurred as previously described (Do-Monte et al., 2013). Only rats with electrode wire tracks that terminated in the target area were included in statistical analyses.

### Statistics and reproducibility

Digital video cameras (Logitech C920) were used to capture rat activity and indices of behavior were obtained using automated video-tracking software (ANY-maze, Stoelting). Handscoring was performed by an experimenter blind to the animal group identity and phase of the behavioral session. All graphic data are presented as mean ± standard error of the mean (s.e.m.). All experiments were replicated in 14 separate batches, with the exception of the homecage opioid administration test, which was performed on a single cohort of rats. After histological verification of correct anatomical location of electrodes, data were combined to generate each group’s sample size. Shapiro–Wilk normality test was performed before all the statistical analyses to determine parametrical or non-parametrical statistical tests. For normal data, we applied unpaired Welch’s t-test or paired Student’s t-test, repeated-measures analysis of variance (followed by Šidák’s post hoc comparison), Z test, Fisher’s Exact test, or Chi-square test, whereas for non-parametric data, we used Mann-Whitney U test as indicated for each experiment using GraphPad Prism 9 software. The sample size was based on estimations by power analysis with a level of significance of 0.05 and a power of 0.8.

#### K-means clustering of behavioral data

K-means cluster analysis was employed to categorize behavioral phenotypes among morphine-treated rats (10 mg/kg) based on their responses in a set of behavioral tests. Data for this analysis were compiled from three measures: freezing (% time in seconds spent freezing during the conflict test), place preference (% change from baseline in time in seconds spent in the drug-paired side during the preference test), and cat odor aversion (% change from preference test in time in seconds spent in the drug/cat-paired side during the conflict test). The number of clusters for the k-means analysis was determined using the elbow method (Thorndike, 1953). This method involved visual inspection of an elbow graph plotting the sum of squared distances against the number of clusters. The chosen number of clusters was based on the point where the graph showed a marked bend, indicating an optimal balance between the number of clusters and the variance explained. The sum of squared distances for 1, 2, and 3 clusters were 16.71, 9.87, and 8.45, respectively, with two clusters being selected as the optimal number for this analysis. The k-means analysis was conducted using DATAtab, an online statistical calculator, with a setting of 10 repetitions for the algorithm (DATAtab Team, 2023). This setting determines the number of times the k-means process is run with different centroid seeds, aiming to provide a more reliable result by minimizing the effect of random initial centroid placement. The results of the cluster analysis were visualized using a 3D scatter plot, representing each rat’s data across the three measured behaviors (see Fig. 2A and Supplementary Fig. 5A). This visualization aided in interpreting the two clusters as indicative of two distinct behavioral phenotypes among the subgroups of rats. For exploring overlap of saline group data with Risk-Avoider and Risk-Taker data in 3D cluster space, group minimum and maximum values for the three behavioral measures were designated as criteria for inclusion in a cluster. Thus, individual rats were considered as being included in a cluster only if their behavioral values fell within the cluster’s minimum and maximum values for those behaviors.

#### Spatial decoding of drug and threat contexts via electrophysiological dissection

Our investigation into risk-taking populations involves a meticulous analysis of electrophysiological data obtained through PL single-unit recordings, coupled with behavioral observations captured in videos from morphine-conditioned place preference and approach-avoidance conflict tests. Recording sessions, each lasting approximately 10 minutes, were conducted at a rate of approximately 15 frames per second, documenting the rats’ task performance.

The spatial representation analysis pipeline began with extracting the rat’s head coordinates from recorded video sessions using the pre-trained machine learning model, DeepLabCut (Mathis et al., 2018). Single-unit electrophysiological data and head positions were then preprocessed, combined, and analyzed to generate heatmaps, z-score matrices, and classify neurons based on responsiveness. Custom Python code for this analysis is available on GitHub (https://github.com/Domontelab/Quave_et_al_2024).

#### Head location extraction

DeepLabCut was used to determine the rat’s head location in each video frame. Frames with a likelihood score below 0.8 were discarded.

#### Data Preprocessing

1. Grid calculation: The spatial layout of the apparatus was divided into bins for spatial analysis. A manually selected square on the arena floor served as a reference for creating the grid, accounting for differences in camera positions across sessions. These coordinates were saved as a ‘.pkl’ file, defining the size and number of bins along the x- and y-axes.
2. Duration alignment: Video durations were standardized to 10 minutes, either by truncating or interpolating frames, ensuring temporal synchronization between behavioral and electrophysiological data.
3. Temporal transformation: Neuronal recording timestamps (1ms resolution) were converted to video frame time by averaging spike counts within each frame interval, resulting in a list of accumulated spikes per frame.

#### Spatial mapping and analysis

1. Heatmap calculation: Using the previously defined spatial grid, heatmaps were generated to represent the pixel coverage of the rat’s head in each spatial bin. The analysis distinguished the paired side, located on the right.
2. Z-score matrix calculation: Time spent in each spatial bin was calculated as the total duration that the rat’s head was present in the bin. The firing rate (Hz) per spatial bin was computed as the accumulated spike count in each spatial bin. The Z-score matrix was then calculated using a time window of 6 seconds per temporal bin.

Classification by neuronal responsiveness: Neurons were classified as responsive based on Z-score thresholds: -1.96 for inhibitory responses, and 2.58 for excitatory responses. Group-specific excitatory and inhibitory Z-scores for responsive neurons were summed, visualized as heatmaps, and saved as ‘.pkl’ files for Risk-Takers, Risk-Avoiders, and saline-treated rats.

#### Spectral clustering of single-unit firing rate data

Spectral clustering analysis of PL neuronal firing rate data was conducted using modified codes from Namboodiri et al., 2019 (Namboodiri et al., 2019), available on GitHub (https://github.com/Domontelab/Quave_et_al_2024). Specifically, PL neuron activity was aligned to the rat’s head transition from the neutral side to the paired side of the CPP apparatus during preference or conflict tests and involved calculating the average firing rates of neurons across all such transitions within a 3-second timeframe before and after in 100 ms bins. These firing rate data were then transformed into Z-scores using the mean and standard deviation from the -3s to -1s pre-transition period. The Z-scored data from preference and conflict tests were merged to form a 120-dimensional dataset per neuron. Principal component analysis (PCA) was employed to simplify these complex data, with the number of principal components for projection into a lower-dimensional space determined by the elbow method. The spectral clustering technique in the Scikit-learn package, utilizing an affinity matrix built from a k-nearest neighbor connectivity matrix, was applied for the clustering process. Several nearest neighbor counts (10, 50, 100, 200, 300) were tested. The optimal parameters and cluster numbers were identified by maximizing the silhouette score in a comprehensive parameter grid search. The separation between clusters was verified by tSNE transformation and visualized in 2D space of tSNE scores between each pair of the clusters against each other. Post-clustering, each neuron received an appropriate cluster label.

---

**Supplementary Figure 1.**
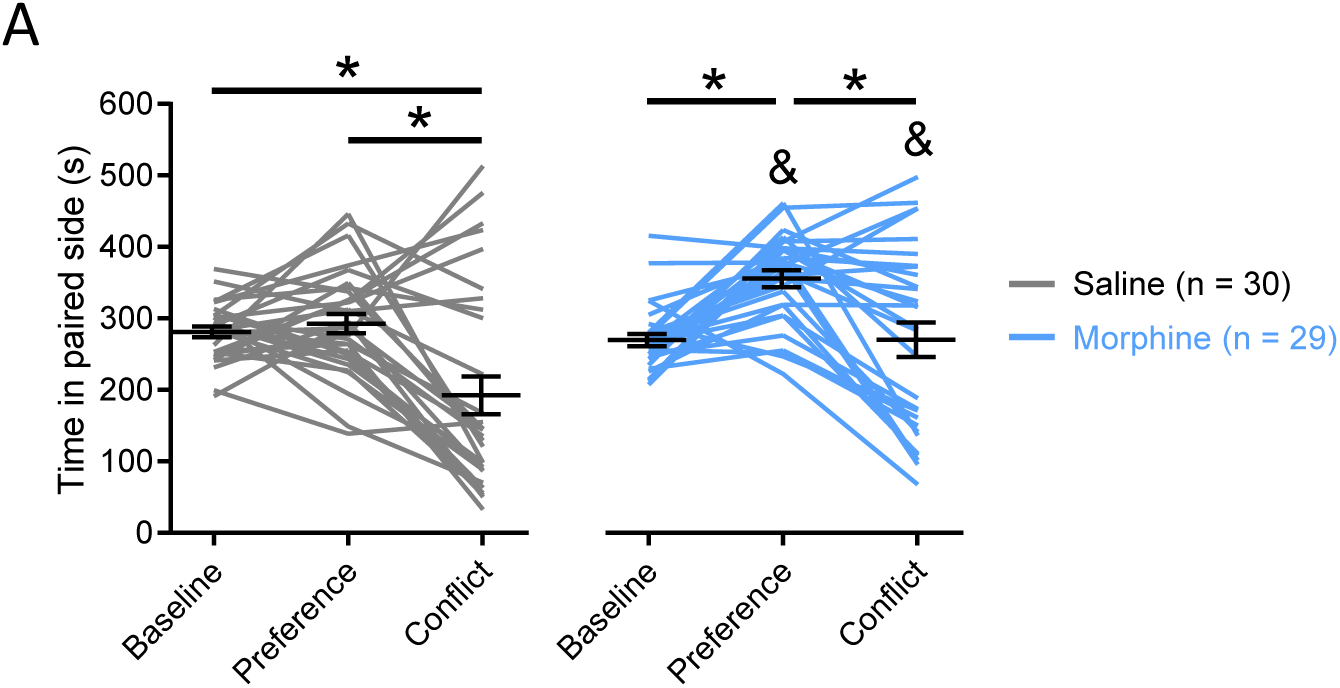
Raw paired side times during the Preference and Conflict Tests for saline- and morphine-treated rats. **A)** Times spent in the paired side during baseline, preference, and conflict tests. Saline-treated rats (gray lines) spent less time in the paired side of the apparatus during the Conflict Test than during either baseline (*p <* 0.001) or the preference test (*p <* 0.0001). Additionally, Morphine-treated rats (blue lines) showed increased time spent in the paired side during the preference test as compared to baseline (*p<* 0.0001) or to Saline-treated rats during the preference test (*p=* 0.029). During the conflict test, Morphine group rats spent less time in the paired side than during the preference test (*p* < 0.0001), but more time in the paired side than Saline rats did during the conflict test (*p* = 0.0043). All observed statistical effects were the results of Šidák’s multiple comparisons tests after two-way repeated measures ANOVA (F (2, 114) = 4.990, Day X Drug interaction, *p* = 0.0084). Ampersand (&) denotes group difference within same test phase. Data are shown as mean ± SEM.

**Supplementary Figure 2.**
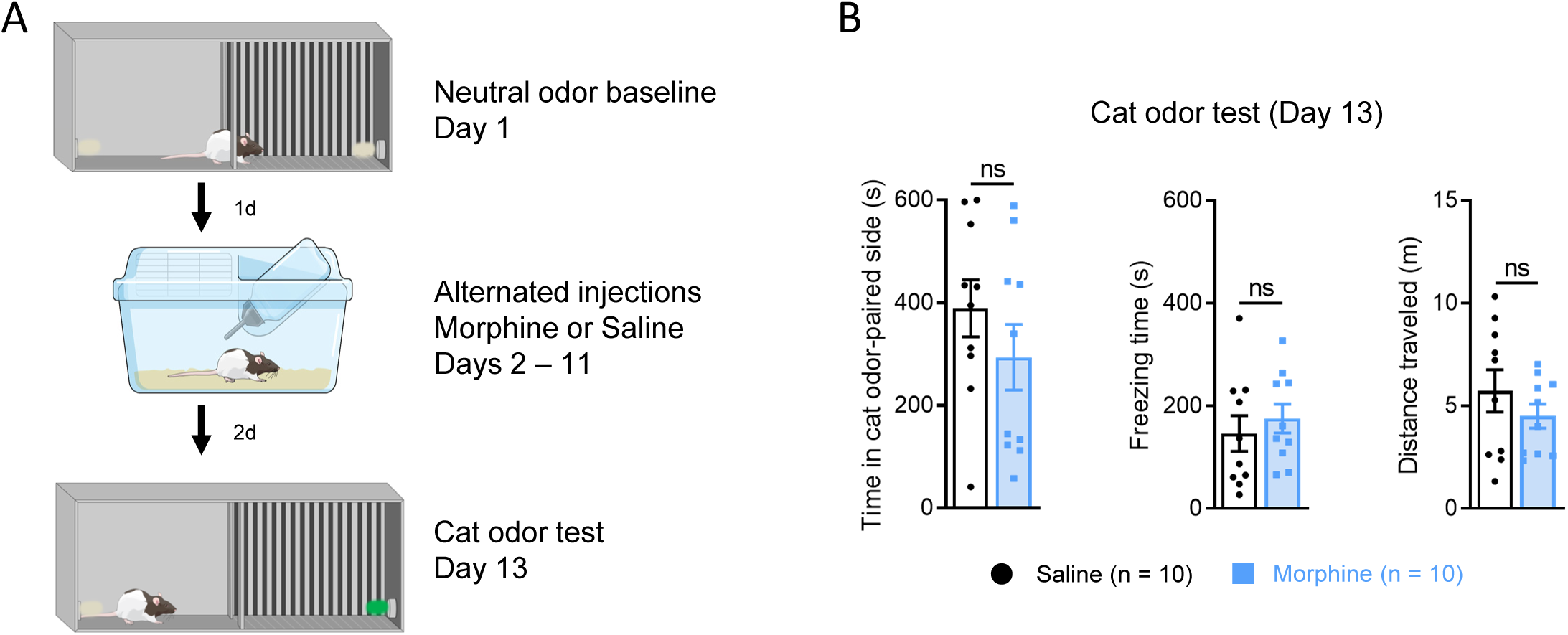
Repeated morphine administration in the home cage does not affect defensive behavior. **A)** Schematic showing experimental timeline of home cage drug administration and cat odor testing. No differences were observed between saline- and morphine-treated rats in **B *left*)** time spent in the cat odor-paired side (Welch’s t-test, *p* = 0.27), **B *center*)** total time spent freezing (Welch’s t-test, *p* = 0.528), or **B *right*)** total distance traveled (Welch’s t-test, *p* = 0.318) during the cat odor test. ns = not significant.

**Supplementary Figure 3.**
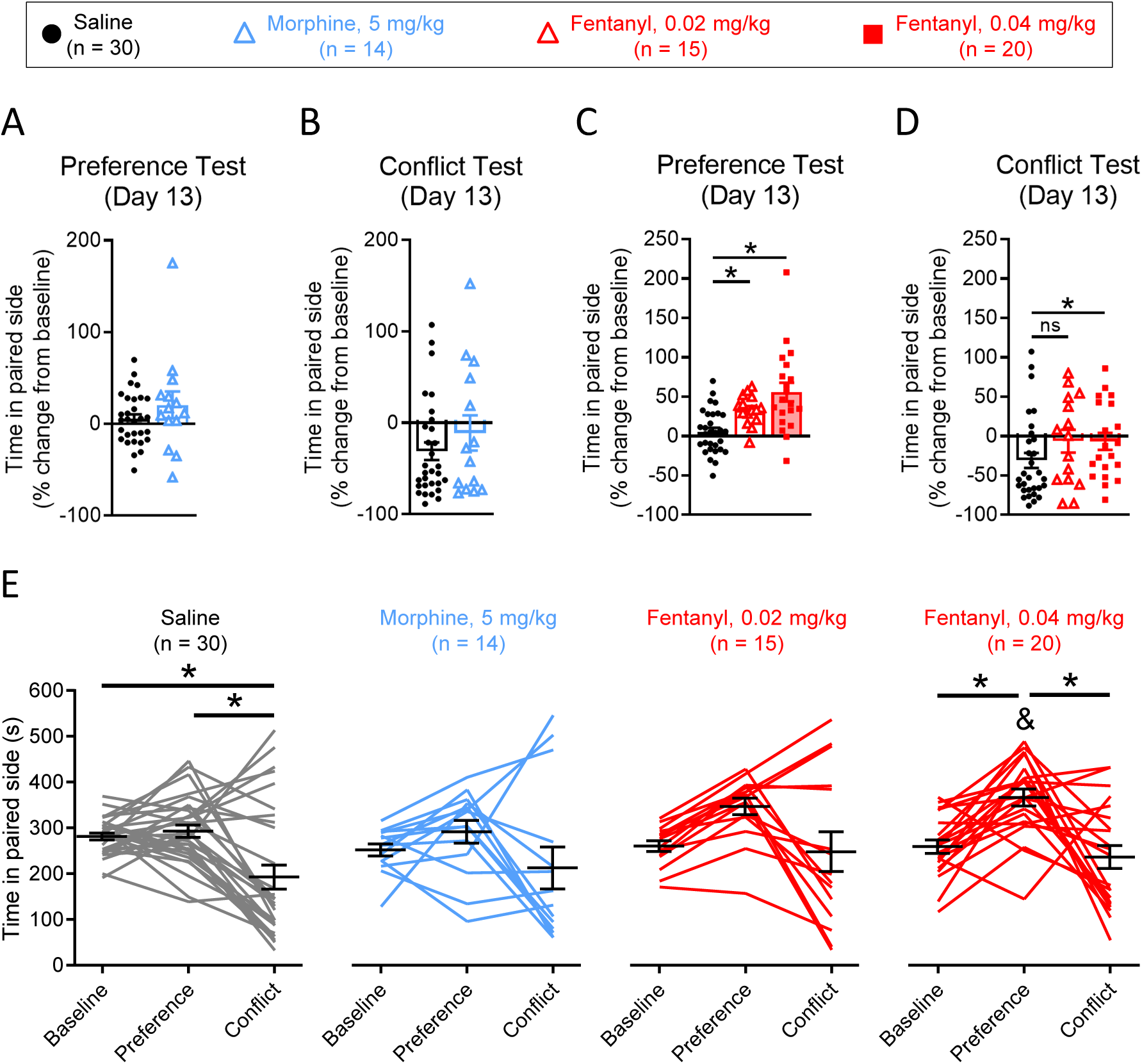
Effects of alternate doses of morphine or fentanyl on male rat behavior. **A)** Percentage of change from baseline in time spent in the drug-paired side of the apparatus by morphine-treated rats during the preference test. 5 mg/kg morphine-treated rats did not exhibit conditioned place preference compared to saline controls (Shapiro-Wilk normality test, *p* = 0.035, Mann-Whitney *U*-test, *p* = 0.336). **B)** Percentage of change from baseline in time spent in the drug/cat-paired side of the apparatus by morphine-treated rats during the conflict test. Rats conditioned with 5 mg/kg morphine showed no difference in aversion to the drug/cat-paired side as compared to saline-treated rats (Shapiro-Wilk normality test, both *p* values < 0.05, Mann-Whitney *U*-test, *p* = 0.54). **C)** Percentage of change from baseline in time spent in the drug-paired side of the apparatus by fentanyl-treated rats during the preference test. Rats treated with either dose of fentanyl exhibited conditioned place preference as measured by the increased amount of time in the drug-paired side compared to saline controls (0.02 mg/kg: Welch’s t-test, *p* < 0.001; 0.04 mg/kg: Welch’s t-test, all *p* values < 0.001). **D)** Percentage of change from baseline in time spent in the drug/cat-paired side of the apparatus by fentanyl-treated rats during the conflict test. Rats conditioned with either saline or 0.02 mg/kg fentanyl showed aversion to the drug/cat-paired side during the conflict test (Shapiro-Wilk normality test, *p* < 0.001, Mann-Whitney test, *p* = 0.14), whereas rats treated with 0.04 mg/kg fentanyl did not (Shapiro-Wilk normality test, *p* < 0.001, Mann-Whitney *U*-test, *p* = 0.0396). **E)** Times spent in the paired side during baseline, preference, and conflict tests by saline-treated rats (gray lines), 5 mg/kg morphine-treated rats (blue lines), 0.02 mg/kg fentanyl-treated rats (red lines, *left*), and 0.04 mg/kg fentanyl-treated rats (red lines, *right*). All statistical tests involved two-way repeated measures ANOVA followed by Šidák’s multiple comparisons tests, each comparing Saline-treated rats to one of three drug-treated groups (Saline vs. 5 mg/kg morphine: Test main effect, F (2, 84) = 9.13, *p* < 0.0001, no Drug x Test interaction, F (2, 84) = 0.66, *p* = 0.52; Saline vs. 0.02 mg/kg fentanyl: Test main effect, F (2, 86) = 12.93, *p* <0.0001, no Drug X Test interaction, F (2, 86) = 2.503, *p* = 0.0878; Saline vs. 0.04 mg/kg fentanyl: Drug x Test interaction, F (2, 98) = 3.704, *p* = 0.0281). Saline-treated rats (gray lines) showed aversion to the drug/cat paired side during the conflict test, as compared to baseline (*p* = 0.0006) or the preference test (*p<* 0.0001). 5 mg/kg morphine-treated rats (blue lines) did not show differences across days, and no group differences were observed when compared to saline-treated rats (gray lines). Similarly, no group differences were observed between saline-treated rats (gray lines) and 0.02 mg/kg fentanyl treated rats (red lines, *left*). 0.04 mg/kg fentanyl-treated rats (red lines, *right*) showed increased time spent in the paired side during the preference test as compared to baseline (*p <* 0.0001) or to saline-treated rats (gray lines) during the preference test (*p =* 0.021). During the conflict test, 0.04 mg/kg fentanyl-treated rats (red lines, right) spent less time in the paired side than during the preference test (*p* < 0.0001), but not when compared to baseline (*p=* 0.792). Ampersand (&) denotes group difference from saline-treated rats within the same test phase. Data are shown as mean ± SEM.

**Supplementary Figure 4.**
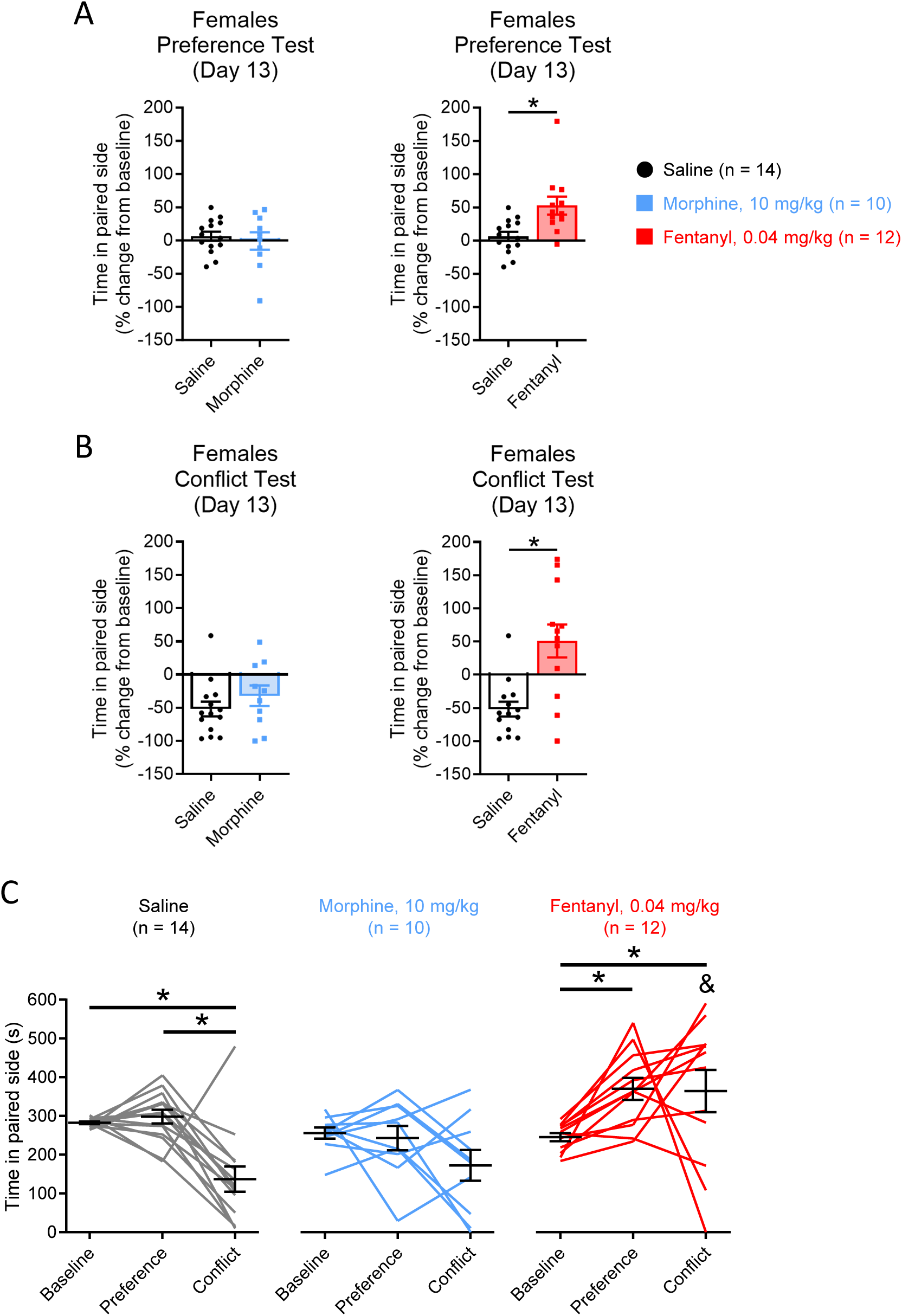
Effects of conditioning with morphine or fentanyl on behavior in female rats. **A *left*)** Percentage of change from baseline in time spent in the drug-paired side of the apparatus by morphine-treated female rats during the preference test. 10 mg/kg morphine-treated rats did not exhibit conditioned place preference compared to saline controls (Welch’s t-test, *p* = 0.64). **A *right*)** Percentage of change from baseline in time spent in the drug-paired side of the apparatus by fentanyl-treated female rats during the preference test. Rats treated with 0.04 mg/kg fentanyl exhibited conditioned place preference as measured by the increased amount of time in the drug-paired side compared to saline controls (Shapiro-Wilk normality test, *p* = 0.017, Mann-Whitney *U*-test, *p* = 0.0011). **B *left*)** Percentage of change from baseline in time spent in the drug/cat-paired side of the apparatus by morphine-treated female rats during the conflict test. Rats conditioned with 10 mg/kg morphine showed no difference in aversion to the drug/cat-paired side as compared to saline-treated rats (Shapiro-Wilk normality test, *p* = 0.044, Mann-Whitney *U*-test, *p* = 0.37). **B *right*)** Percentage of change from baseline in time spent in the drug/cat-paired side of the apparatus by fentanyl-treated female rats during the conflict test. Rats conditioned with saline showed aversion to the drug/cat-paired side during the conflict test, whereas rats treated with 0.04 mg/kg fentanyl did not (Shapiro-Wilk normality test, *p* = 0.044, Mann-Whitney *U*-test, *p* = 0.0031). **C)** Times spent in the paired side during baseline, preference, and conflict tests by female saline-treated rats (gray lines), 10 mg/kg morphine-treated rats (blue lines), and 0.04 mg/kg fentanyl-treated rats (red lines). All statistical tests involved two-way repeated measures ANOVA followed by Šidák’s multiple comparisons tests, each comparing Saline-treated rats to one of the two drug-treated groups (Saline vs. 10 mg/kg morphine: Test main effect, F (2, 44) = 13.23, *p* < 0.0001, no Drug x Test interaction, F (2, 44) = 1.62, *p* = 0.209; Saline vs. 0.04 mg/kg fentanyl: Test main effect, F (2, 48) = 4.39, *p* = 0.018, Drug X Test interaction, F (2, 48) = 9.6, *p* = 0.0003). Both saline-treated rats (gray lines) and morphine-treated rats (blue lines) showed aversion to the drug/cat-paired side during the conflict test (baseline vs. conflict, *p* = 0.0074); no group differences were observed. Fentanyl-treated rats (red lines) showed increased time spent in the paired side during both the preference test (*p=* 0.022) and the conflict test as compared to baseline (*p* = 0.031). Fentanyl-treated rats (red lines) also spent more time in the paired side than saline-treated rats did during the conflict test (*p* < 0.0001). Ampersand (&) denotes group difference from saline-treated rats within the same test phase. Data are shown as mean ± SEM.

**Supplementary Figure 5.**
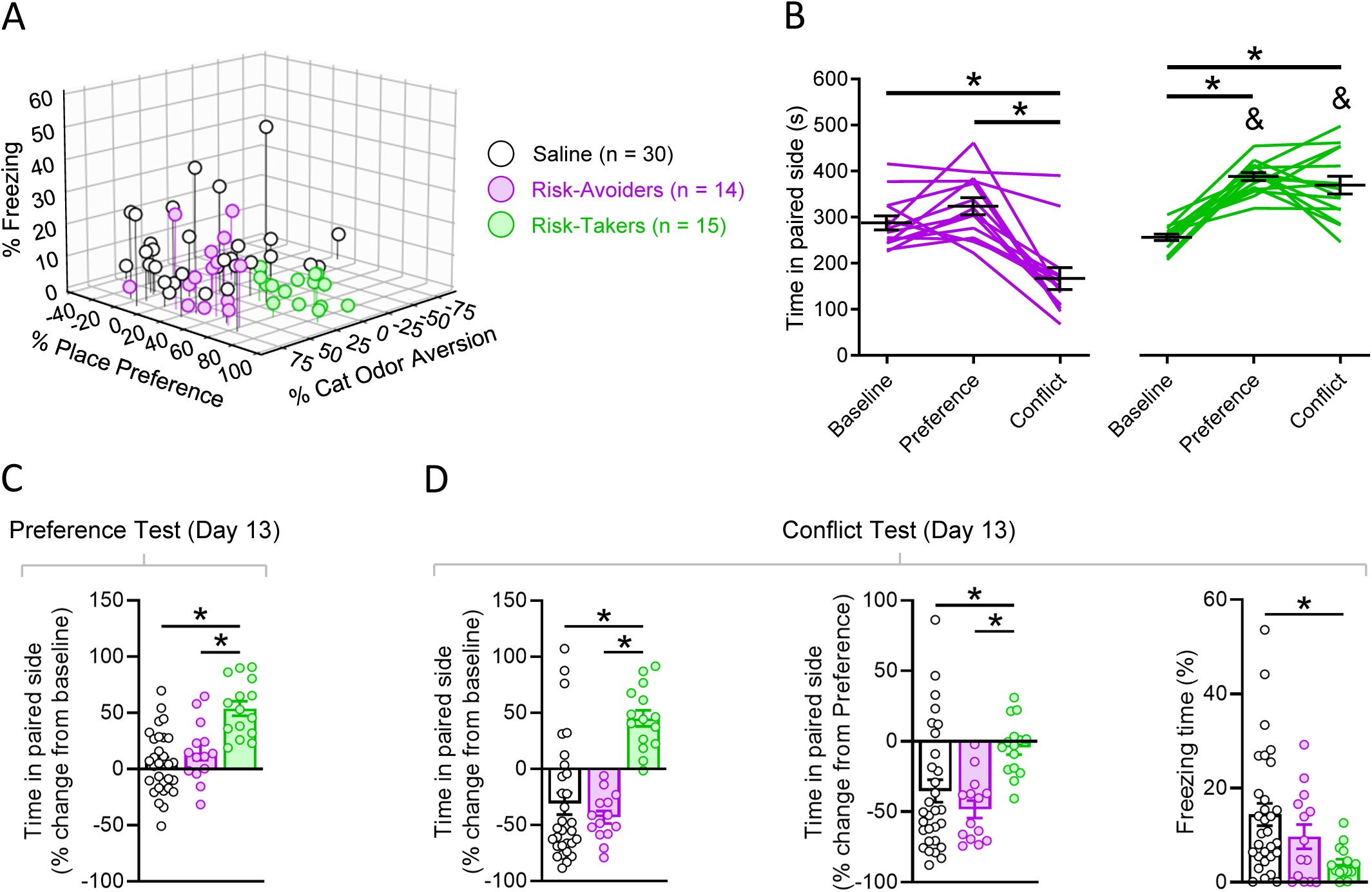
Effects of morphine conditioning on Risk-Takers and Risk-Avoiders compared to saline-treated rats. **A)** 3-dimensional plot of data from saline-treated rats (black/white cluster, n = 30), Risk-Avoiders (magenta cluster, n = 15), and Risk-Takers (green cluster, n = 14) on measures of freezing (% time spent freezing during the conflict test), place preference (% change from baseline in time spent in the drug-paired side during the preference test), and cat odor aversion (% change from preference test in time spent in the drug/cat-paired side during the conflict test). **B)** Times spent in the paired side during baseline, preference, and conflict tests. Risk-Avoiders (magenta lines) spent less time in the paired side of the apparatus during the conflict test than during either baseline (*p <* 0.0001) or the preference test (*p <* 0.0001). Risk-Takers (green lines) showed increased time spent in the paired side during the preference test as compared to baseline (*p <* 0.0001), and as compared to Risk-Avoiders during the preference test (*p* = 0.019). During the conflict test, Risk-Takers spent more time in the paired side than at baseline (*p <* 0.0001), and as compared to Risk-Avoiders during the conflict test (*p* < 0.0001; two-way repeated measures ANOVA, Group x Test interaction, F (2, 54) 37.93, *p <* 0.0001, Šidák’s multiple comparisons tests). Ampersand (&) denotes group difference within same test phase. **C)** Percentage of change from baseline in time spent in the drug-paired side of the apparatus. Risk-Takers demonstrated greater place preference than either saline-treated rats (*p* < 0.0001) or Risk-Avoiders (*p* = 0.0008). Saline-treated rats did not differ from Risk-Avoiders on preference measures (*p* = 0.62; one-way ANOVA, F (2, 56) = 16.69, *p* < 0.0001, Šidák’s multiple comparisons tests). **D *left*)** Percentage of change from baseline in time spent in the drug/cat-paired side of the apparatus during the conflict test. Risk-Takers showed less cat odor aversion than either saline-treated rats (*p* < 0.0001) or Risk-Avoiders (*p* < 0.0001). Saline-treated rats did not differ from Risk-Avoiders on this measure (*p* = 0.75; one-way ANOVA, F (2, 56) = 20.67, *p* < 0.0001, Šidák’s multiple comparisons tests). **D *center*)** Percentage of change from the preference test in time spent in the drug/cat-paired side of the apparatus during the conflict test. Risk-Takers showed less cat odor aversion than either saline-treated rats (*p* = 0.031) or Risk-Avoiders (*p* = 0.0039). Saline-treated rats did not differ from Risk-Avoiders on this measure (*p* = 0.59; one-way ANOVA, F (2, 56) = 6.302, *p* = 0.0034, Šidák’s multiple comparisons tests). **D *right*)** Percentage of time spent freezing during the conflict test. Risk-Takers and Risk-Avoiders displayed similar levels of freezing during the conflict test (*p* = 0.39). However, Risk-Takers showed decreased levels of freezing during the conflict test when compared to saline-treated rats (*p* = 0.0089). Saline-treated rats did not differ from Risk-Avoiders on this measure (*p* = 0.44; one-way ANOVA, F (2, 56) = 4.9, *p* = 0.011, Šidák’s multiple comparisons tests). Data are shown as mean ± SEM.

**Supplementary Figure 6.**
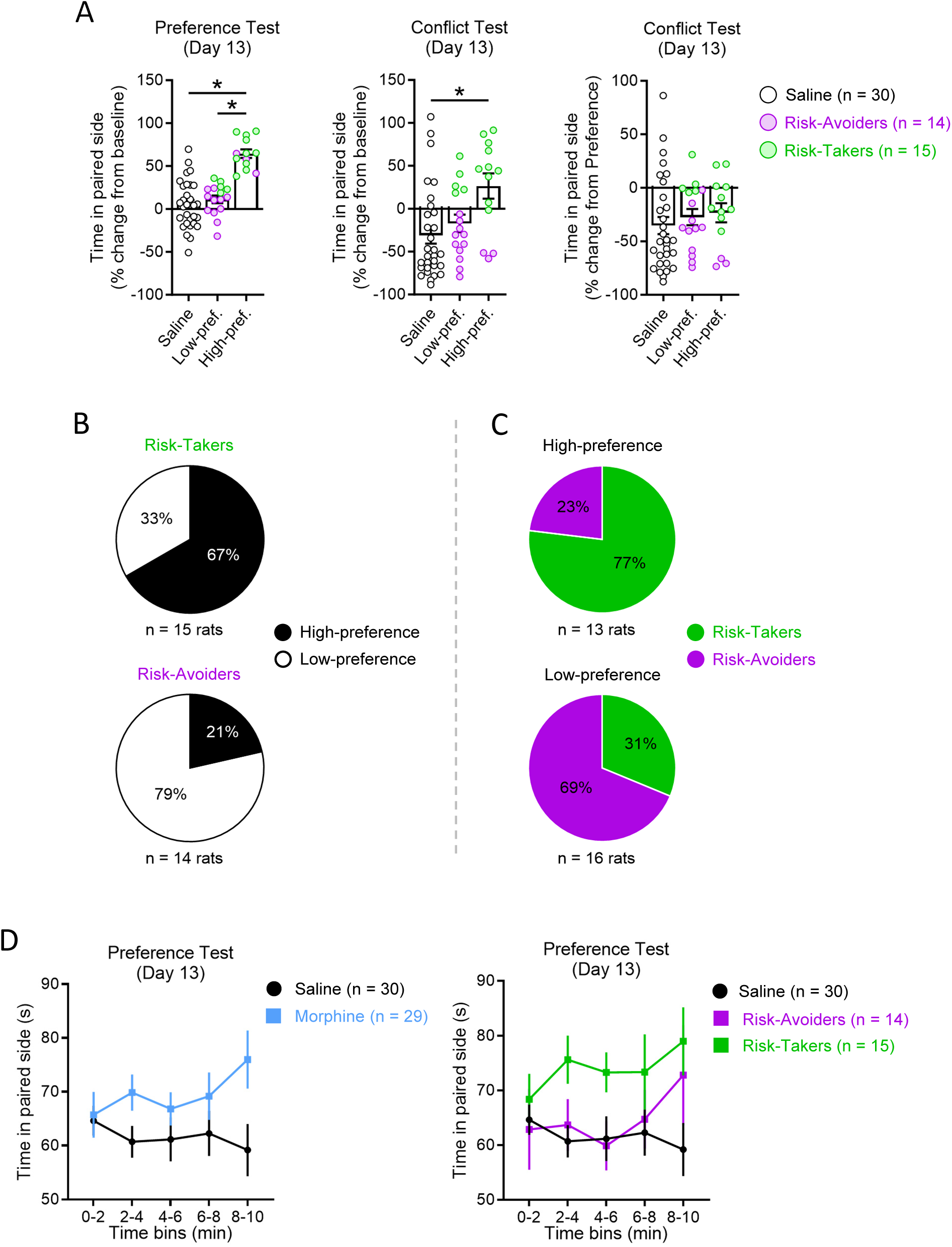
Investigations of extinction, strength, and maintenance of CPP across rat subgroups. **A *left*)** Percentage of change from baseline in time spent in the drug-paired side of the apparatus during the preference test. High-preference rats demonstrated greater place preference than either saline-treated rats (*p* < 0.0001) or low-preference rats (*p* < 0.0001). Saline-treated rats did not differ from low-preference rats on preference measures (*p* = 0.80; one-way ANOVA, F (2, 56) = 29.96, *p* < 0.0001, Šidák’s multiple comparisons tests). **A *center*)** Percentage of change from baseline in time spent in the drug/cat-paired side of the apparatus during the conflict test. High-preference rats showed less cat odor aversion than saline-treated rats (*p* = 0.0032) but not low-preference rats (*p* = 0.0697). Saline-treated rats did not differ from low-preference rats on this measure (*p* = 0.76; one-way ANOVA, F (2, 56) = 5.968, *p* = 0.0045, Šidák’s multiple comparisons tests). **A *right*)** Percentage of change from the preference test in time spent in the drug/cat-paired side of the apparatus during the conflict test. High-preference rats did not show differences in cat odor aversion from either saline-treated rats (*p* = 0.72) or low-preference rats (*p* = 0.99), nor did saline-treated rats did not differ from low-preference rats on this measure (*p* = 0.88; one-way ANOVA, F (2, 56) = 0.5207, *p* = 0.597, Šidák’s multiple comparisons tests). **B)** Percentages of Risk-Takers and Risk-Avoiders that were also classified as either high-preference or low-preference rats. The Risk-Taker group was comprised of a greater percentage high-preference rats than was the Risk-Avoider group (Fisher’s Exact test, *p* = 0.025). **C)** Percentages of high-preference and low-preference rats that were also classified as either Risk-Takers or Risk-Avoiders. The high-preference group was comprised of a greater percentage of Risk-Takers than was the low-preference group (Fisher’s Exact test, *p* = 0.025). **D)** Time spent in the drug-paired side of the apparatus across time (2-min bins) during the preference test. No group showed a change in time spent in the drug-paired side across the duration of the preference; this was true for saline-treated and morphine-treated groups (*left*), as well as for Risk-Avoider and Risk-Taker subgroups (*right*). Data represented in line or bar graphs are shown as mean ± SEM.

**Supplementary Figure 7.**
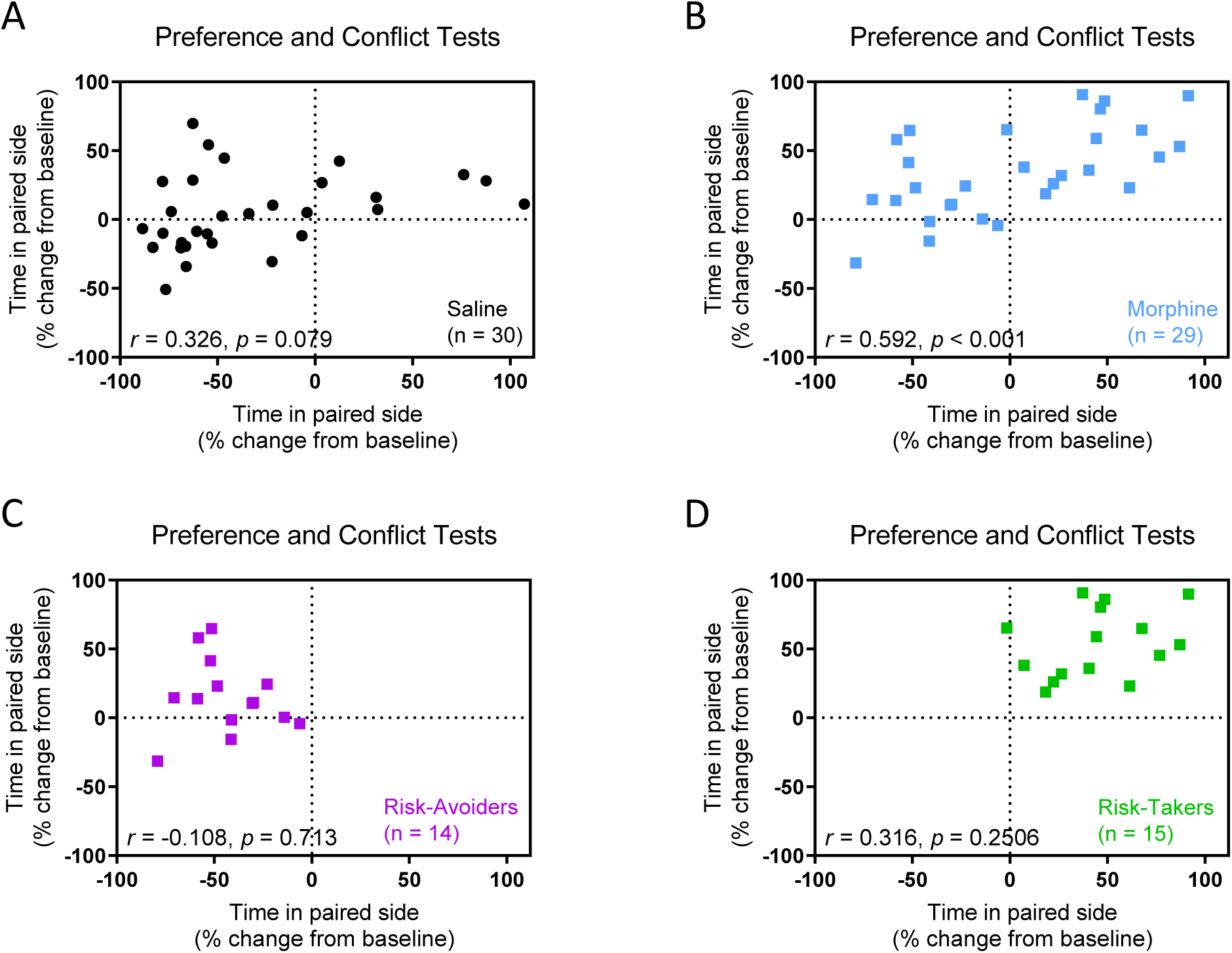
Correlation analyses of place preference and risk-taking behaviors in male morphine-treated rats. **A–D)** Scatterplots of data for analyses of correlation between place preference score (% change from baseline in time spent in the drug paired side of the apparatus during the preference test; y-axes) and risk-taking behavior during conflict (% change from baseline in time spent in the drug paired side of the apparatus during the conflict test; x-axes). While no correlations between place preference score and risk-taking behavior were found in A) saline-treated rats (*r*(28) = 0.33, *p* = 0.08), or within either C) Risk-Avoider (*r*(12) = -0.11, *p* = 0.71) or D) Risk-Taker groups (*r*(13) = 0.32, *p* = 0.25), B) it was observed that place preference scores were positively correlated with risk-taking behavior when data from all morphine-treated rats were analyzed together (*r*(27) = 0.59, *p* < 0.001).

**Supplementary Figure 8.**
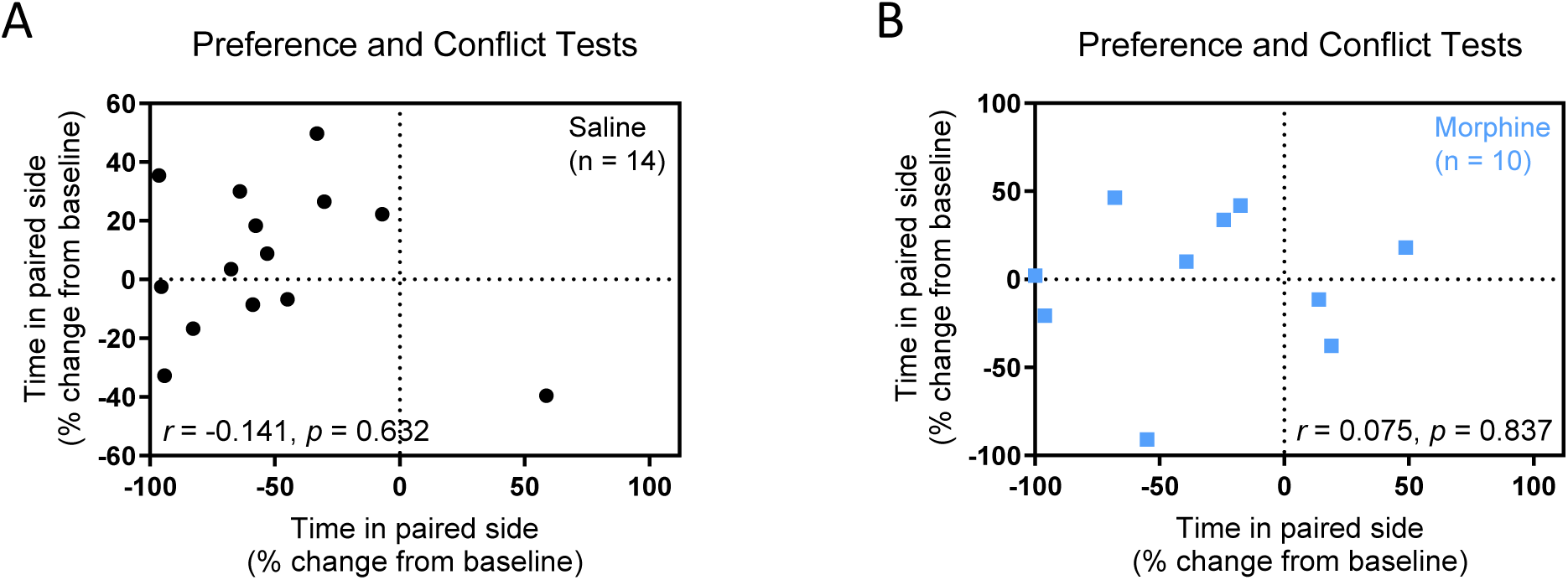
Correlation analyses of place preference and risk-taking behaviors in female morphine-treated rats. **A–B)** Scatterplots of data for analyses of correlation between place preference score (% change from baseline in time spent in the drug paired side of the apparatus during the preference test; y-axes) and risk-taking behavior during conflict (% change from baseline in time spent in the drug paired side of the apparatus during the conflict test; x-axes). No correlation between place preference score and risk-taking behavior was found for either female A) saline-treated rats or B) morphine-treated rats.

**Supplementary Figure 9.**
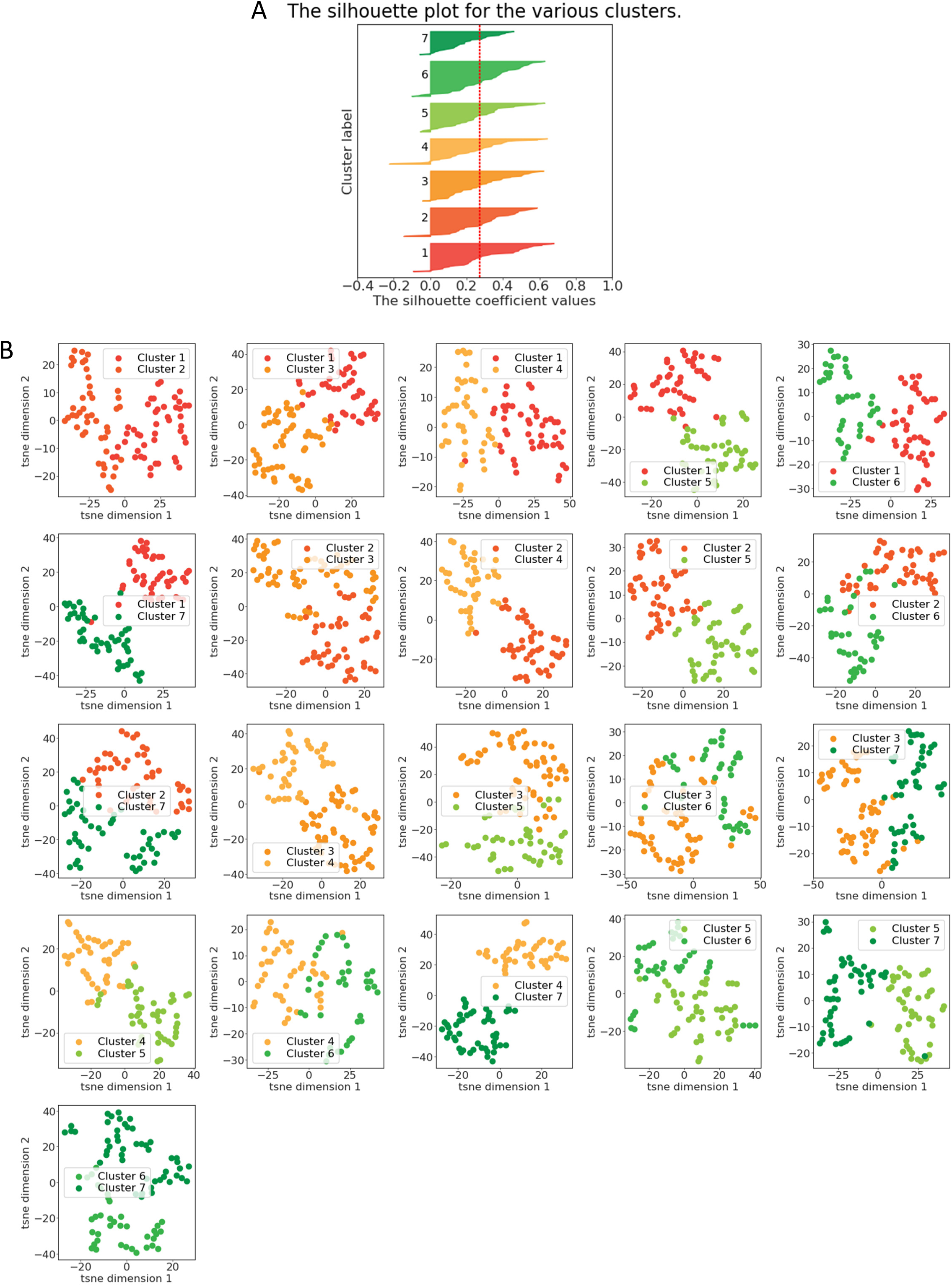
Validation of spectral clustering analysis. **A)** Silhouette score plot for the final clusters. Each row represent the distribution of silhouette scores for each cell in the cluster. Red dashed vertical line indicates the average silhouette score of the clusters, which is the highest across various combination of clustering parameters. B) tSNE plot for each pairs of 7 clusters against each other using the tSNE transformation of the PCA score. As indicated by the plot, most clusters are well separated in this 2D space, suggesting that they are real clusters.

**Supplementary Figure 10.**
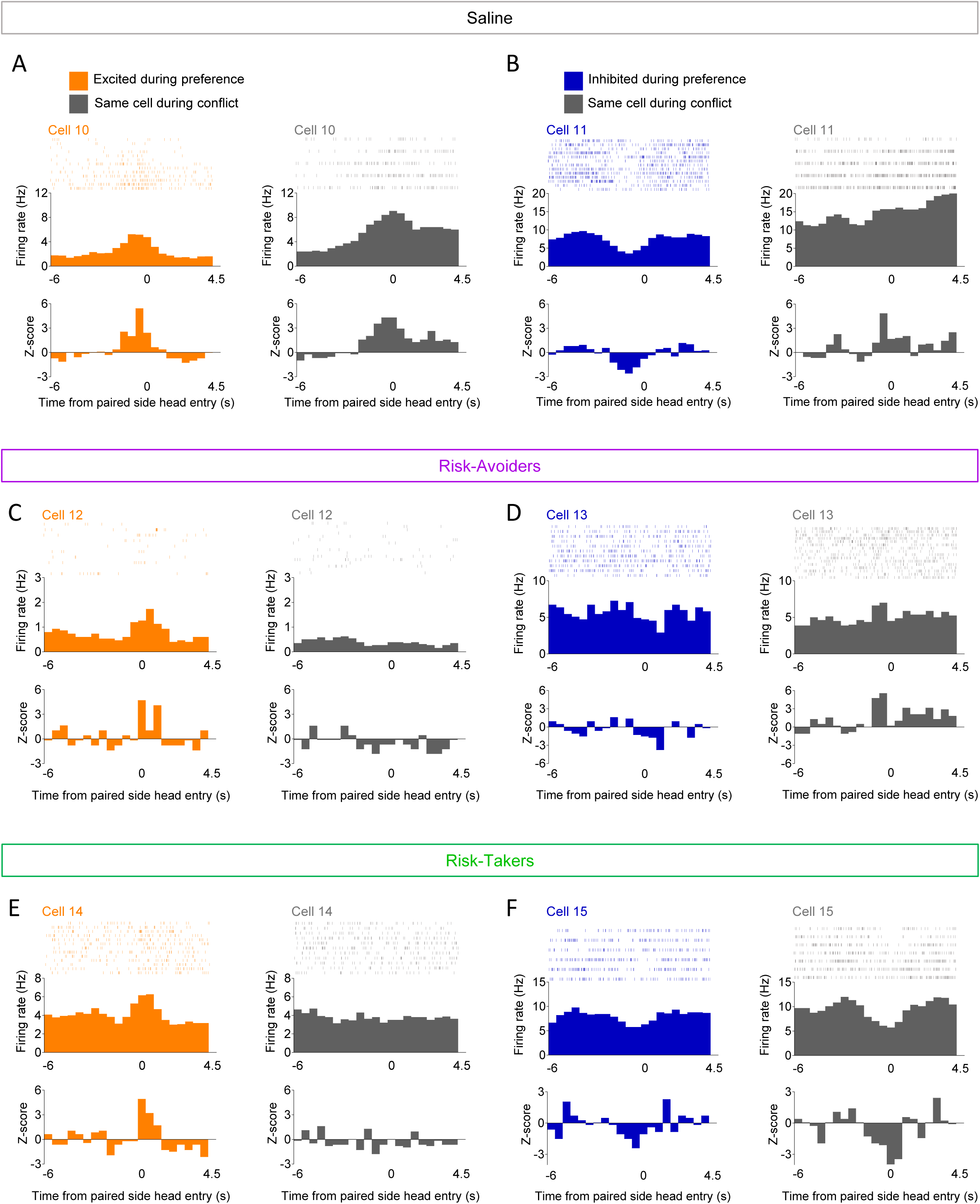
Examples of excitatory and inhibitory single-unit activity during paired-side head entries in the preference and conflict tests. Examples of activity from single cells from **A–B)** a saline treated rat, **C–D)** a Risk-Avoider, and **E–F)** a Risk-Taker aligned to paired-side head entries in the preference and conflict tests. Data are shown as spikes by trial (top of each figure) and peri-stimulus time histograms of data for all trials in firing rate (Hz, middle of each figure) and Z-score (bottom of each figure). Left-side graphs in each figure display the activity of each cell in the preference test, and right-side graphs show activity of the same cell during the conflict test.

## Notes

<u>Conflict of interest statement</u>: The authors declare no competing interests.

### Competing Interest Statement

The authors have declared no competing interest.

### Summary of Updates

We revised the manuscript to clarify previous analyses, incorporate new findings in the Methods and Results sections, and expand the Discussion as requested by reviewers. Additional analyses were conducted to address reviewers' concerns, strengthening our interpretation of the data. We also performed machine learning analyses on behavioral and electrophysiological data to explore the neural correlates of opioid context representation and decision-making during risk-taking behavior. These results are detailed in a new section, which we believe broadens the scope and impact of our study.

